# Moving Beyond Processing and Analysis-Related Variation in Neuroscience

**DOI:** 10.1101/2021.12.01.470790

**Authors:** Xinhui Li, Nathalia Bianchini Esper, Lei Ai, Steve Giavasis, Hecheng Jin, Eric Feczko, Ting Xu, Jon Clucas, Alexandre Franco, Anibal Sólon Heinsfeld, Azeez Adebimpe, Joshua T. Vogelstein, Chao-Gan Yan, Oscar Esteban, Russell A. Poldrack, Cameron Craddock, Damien Fair, Theodore Satterthwaite, Gregory Kiar, Michael P. Milham

## Abstract

When fields lack consensus standard methods and accessible ground truths, reproducibility can be more of an ideal than a reality. Such has been the case for functional neuroimaging, where there exists a sprawling space of tools and processing pipelines. We provide a critical evaluation of the impact of differences across five independently developed minimal preprocessing pipelines for functional MRI. We show that even when handling identical data, inter-pipeline agreement was only moderate, critically shedding light on a factor that limits cross-study reproducibility. We show that low inter-pipeline agreement mainly becomes appreciable when the reliability of the underlying data is high, which is increasingly the case as the field progresses. Crucially, we show that when inter-pipeline agreement is compromised, so too are the consistency of insights from brainwide association studies. We highlight the importance of comparing analytic configurations, as both widely discussed and commonly overlooked decisions can lead to marked variation.

---

As the neuroscience community intensifies its efforts to characterize the neural bases of individual differences in brain and behavior, there has been a growing appreciation of the importance of measurement reliability^1–4^. Theoretical and empirical studies have emphasized reliability as an upper bound for validity^5^, as well as a determinant of statistical power and observable effect sizes^6,7^. This increased focus on quantifying and optimizing reliability is particularly important for studies that use functional magnetic resonance imaging (fMRI), where reliability is an essential prerequisite for clinical translation^8,9^. Specifically, a multitude of studies have pointed to the ability to dramatically improve measurement reliability by increasing the amount of fMRI data obtained per individual (i.e., 25+ minutes vs. the more traditional 5–10 minutes^10^), improved acquisitions (e.g., multiecho fMRI^11^), or robust analytic strategies (e.g., bagging, multivariate modeling^12,13^).

However, multiple forms of reliability exist. Most prior efforts in neuroimaging have focused on test-retest reliability^8,14^, which is a critical prerequisite for any laboratory test that aims to quantify individual differences in a stable trait. Another important form of reliability is inter-rater reliability (or agreement), which can refer to reliability across data acquisition instruments (e.g., MRI scanners), or processing and analytic techniques (e.g., pipelines). Although less commonly evaluated, inter-pipeline agreement (IPA) (i.e., the similarity of derived data generated by independent processing pipelines when handling the same data) is critical, as it ensures the suitability of data for comparison and/or aggregation across studies. IPA is particularly important for fMRI analysis, where independently developed tools perform conceptually similar, though not identical, operations.

The presence of a common set of minimal preprocessing steps is assumed to reduce analytic variability and promote reproducibility. However, a growing number of studies suggest that differences in the implementation of these processing steps or how they are “glued together” can yield notably different outcomes. Studies systematically comparing specific preprocessing steps such as segmentation^15^, motion correction^16^, and registration^17–19^ have reported substantial variation in outputs generated across independently developed packages when applied to the same data. In the analysis of task fMRI data, end-to-end pipelines built using different software packages have been found to produce marked variation in the final results^20–23^. Most recently, seventy teams independently analyzed the same dataset with their preferred preprocessing and statistical analysis methods, and reported inconsistent hypothesis test results^24^. While these findings collectively highlight that analytical variability can have substantial effects on the scientific conclusions of neuroimaging studies, there remains a conspicuous lack of clarity regarding the sources of these differences.

Here, we perform a systematic evaluation, replication, and source localization of differences that emerge across fMRI preprocessing pipelines through the lens of functional connectomics. First, we extended the literature examining pipeline implementation-related variation in fMRI by comparing the results generated using minimal preprocessing in five distinct and commonly used pipelines for functional connectivity analysis — Adolescent Brain Cognitive Development fMRI Pipeline (ABCD-BIDS)^25^, Connectome Computational System (CCS)^26^, Configurable Pipeline for the Analysis of Connectomes default pipeline (C-PAC:Default)^27^, Data Processing Assistant for Resting-State fMRI (DPARSF)^28^ and fMRIPrep Long-Term Support version (fMRIPrep-LTS [volume-based pipeline])^29^. As indicated in **Table 1**, while the minimal processing pipelines are generally aligned with respect to their fundamental steps, the specifics of implementation are notably different. See Supplemental Section S1 for a summary of the conceptual differences across pipeline pairs. Second, we demonstrated the role that pipeline replication can play as a means of exploring analytic variation and assessing the robustness of findings. To this end, we leveraged and extended the flexibility of C-PAC to replicate non-MATLAB dependent minimal processing pipelines (ABCD-BIDS, CCS, fMRIPrep-LTS) in a single platform. Third, we put pipeline-related variation into context with more widely studied sources of variability in the imaging literature — namely scan duration (i.e., quantity of data per subject) and global signal regression^30^. We demonstrated that IPA is an upper bound on the overall generalizability of results that will become increasingly apparent as the field: i) improves data acquisition to optimize test-retest reliability for measurements of individual differences, and ii) makes advances to processing techniques. Finally, we evaluated the origins of differences among pipelines, showing that the specific causes of compromises in IPA can vary depending on the pipelines being examined, and raising cautions about the potential impact that seemingly innocuous decisions can have on IPA (e.g., MNI brain template version and resolution). We provide recommendations for improving IPA as the field continues to pursue the goal of reproducible neuroscience.

**Table 1.**
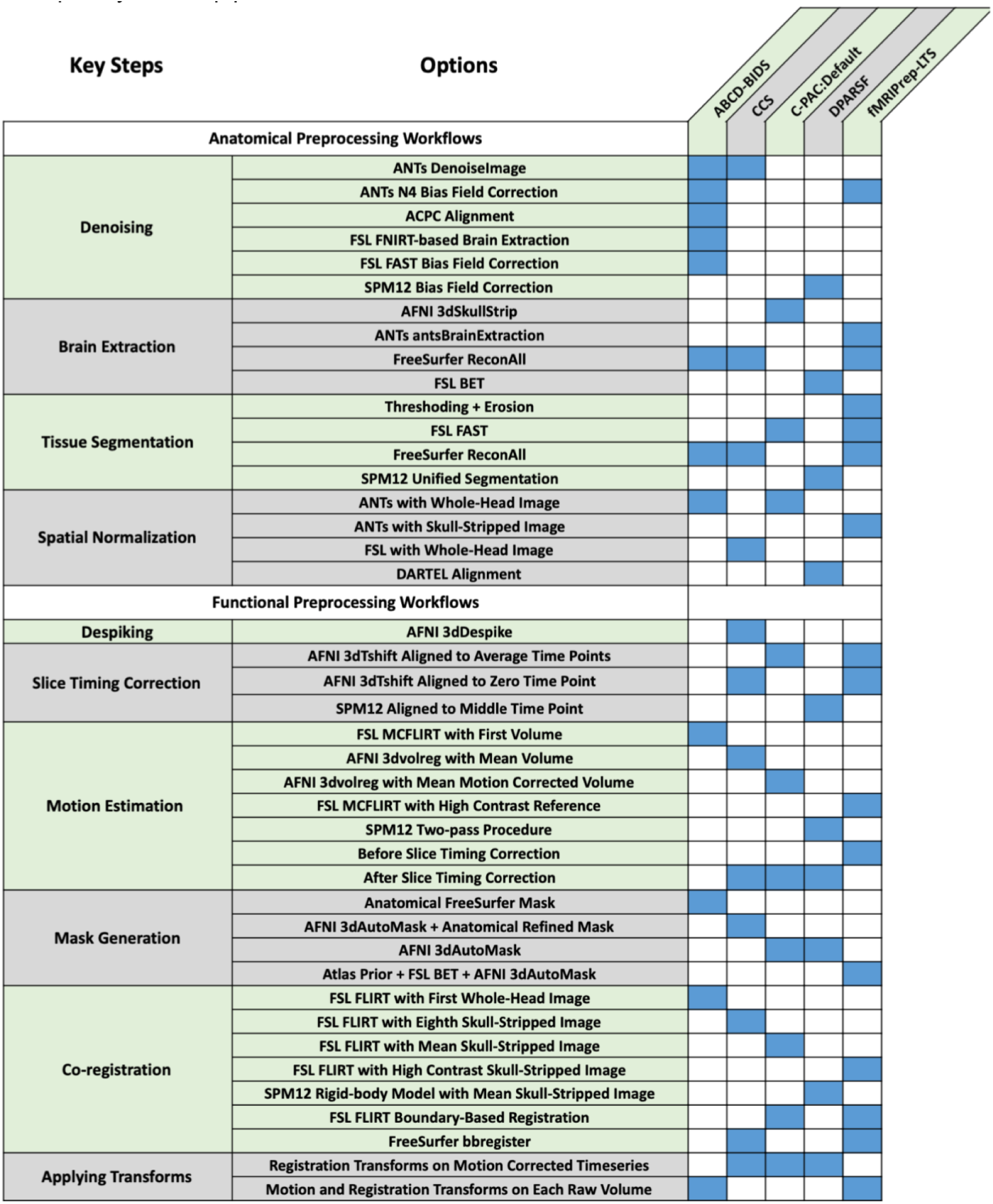
Key methodological differences across five fMRI preprocessing pipelines. For each of the pipeline packages evaluated, rows list libraries and tools used for each processing stage, with blue cells indicating their membership within a pipeline configuration. The heterogeneity across columns illustrates the differences in implementations for even conceptually similar pipelines.

## Results

### Distinct minimal preprocessing pipelines show moderate inter-pipeline agreement

We processed the Hangzhou Normal University (HNU) dataset (29 subjects, 10 sessions, each session has 10-min single-band resting state fMRI per subject, TR = 2000 ms, see Methods for more details), made available through the Consortium of Reliability and Reproducibility (CoRR)^31^. We used each of five different pipelines in widely-used packages for fMRI preprocessing (ABCD-BIDS, CCS, C-PAC:Default, DPARSF, fMRIPrep-LTS). Consistent with prior work^22,24^, we found significant variation in functional connectivity estimates produced using minimally processed data — even when using data from the same session (Kolmogorov-Smirnov test p_corrected_ < 0.001 for all pairs). Findings were robust to the assessment measure (individual-level matrix Pearson correlation, the edge-wise intra-class class correlation coefficient (ICC), the image intraclass correlation coefficient (I2C2)^32^, discriminability^33^) and atlas (Schaefer 200, 600, 1000^34^) used. As depicted in **Figure 1**, among the pipelines, CCS, C-PAC:Default, and fMRIPrep-LTS exhibited the highest degree of IPA with one another, regardless of whether looking at univariate or multivariate perspectives (e.g., Schaefer 200, matrix correlation: 0.811-0.861; ICC: 0.742-0.823; I2C2: 0.785-0.840; discriminability: 1.000). Importantly, across all comparisons, IPA consistently decreased as the dimensionality of the network increased, defined by the number of parcellation units (paired t-test p_corrected_ < 10^-5^ for all pairwise comparisons). The results shown in **Figure 1** can be viewed on the brain surface and within connectivity matrices directly in Supplemental Section S2, and the impact of these variations on downstream extraction of graph theoretic measures or analytic applications can be found in Supplemental Section S3. Specifically, we highlight in our supplementary materials that when handling the same exact data i) pipeline variation leads to significant variation in the sorting of individuals based on graph theoretic measures (e.g., difference in ranks across pipeline pairs was 8.5 of a maximum 14.5 positions on average, demonstrating significant variation in relative individual connectivity). It is important to note that, in all cases, the pipelines compared here do not differ in conceptual approach or design — i.e., global signal regression and other denoising statuses are consistent — such that all variation in both the measures themselves and downstream modeling efforts is reflective of unintended package-related variation.

**Figure 1.**
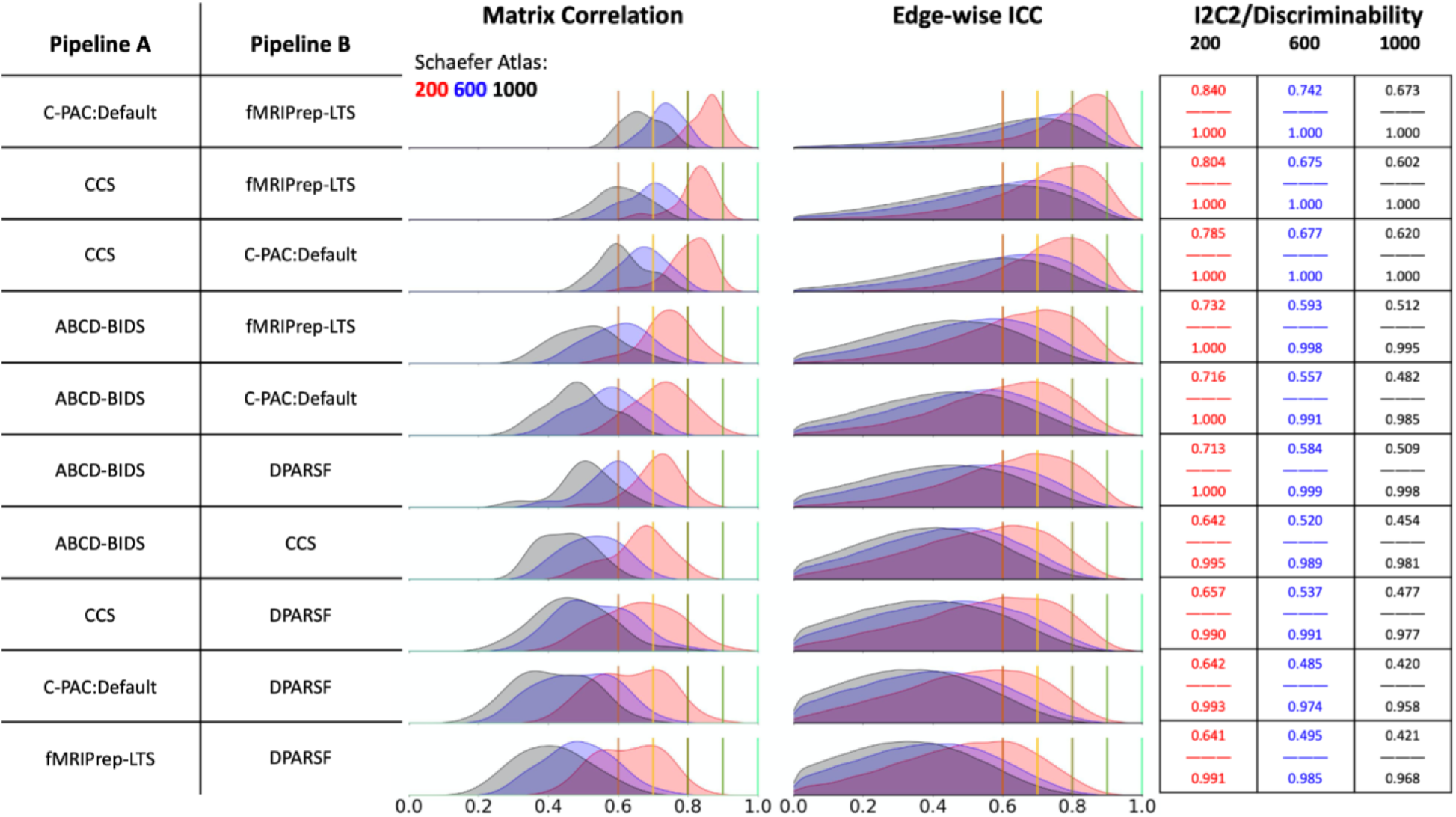
Inter-pipeline agreement for minimal preprocessing in five fMRI preprocessing packages. Each row indicates a pair of pipelines, across which the individual-level matrix Pearson correlation, edge-wise ICC, I2C2, and discriminability were computed for each subject using identical data (i.e., same session). The rows are sorted according to the median matrix Pearson correlation for the Schaefer 200 atlas. Across any pair of pipelines, the median edge-wise ICC — a common measure of result reliability — does not exceed 0.823. Edgewise ICC is also observed to be as low as 0.504, where an accepted reference value for sufficient similarity across raters, in this case pipelines, is typically considered as ICC > 0.9.

In looking at specific packages, DPARSF showed the lowest similarity to the others (e.g., Schaefer 200, matrix correlation: 0.639-0.729; ICC: 0.504-0.612; I2C2: 0.641-0.713; discriminability: 0.990-1.000). This was likely due to the fact that DPARSF was the only SPM/MATLAB-based tool, and uses distinct algorithms, methods, and codebase. ABCD-BIDS, which is based on the HCP Pipelines^35^, showed modest IPA with the other pipelines (e.g., Schaefer 200, matrix correlation: 0.667-0.757; ICC: 0.563-0.651; I2C2: 0.642-0.732; discriminability: 0.995-1.000). This may reflect the fact that ABCD-BIDS is also the most conceptually distinct, including extra denoising and alignment steps for brain extraction. It should also be noted that ABCD-BIDS uniquely doesn’t use boundary-based registration (BBR) unless paired with distortion correction, as prior work suggests that BBR with uncorrected images can lead to misregistration^36^. As discussed later, when we explored sources of variation, repetition of ABCD-BIDS processing using BBR (Supplemental Section S4) yields IPA comparable to that of other non-MATLAB pipelines. To identify the impact of data quality on the above findings, we replicated this exploration using comparably sized low- (n = 29) and high-motion (n = 29) cohorts from the Healthy Brain Network (HBN)^37^, which makes use of the state-of-the-art multiband fMRI sequence from the NIH ABCD Study (5-min resting state fMRI per subject, TR = 800 ms; Supplemental Section S5). We found that, consistent with our findings with the single band HNU sample, the two HBN cohorts showed low levels of IPA (e.g., Schaefer 200, average ICC = 0.832), with IPA only being slightly higher for the low-motion cohort.

### Replicated minimal preprocessing pipelines achieve high inter-pipeline agreement

We investigated minimal preprocessing pipeline differences (**Table 1**) and expanded the configurable options in C-PAC to generate minimal preprocessing pipelines harmonized to match each of the three additional non-MATLAB-based pipelines (ABCD-BIDS, CCS, fMRIPrep-LTS; see Methods for details). A primary goal of this process was to improve the IPA across pipelines to commonly accepted standards (i.e., ICC > 0.9)^38^. **Figure 2** shows the outcome of the pipeline replication process in C-PAC, and demonstrates that median ICC values exceed 0.98 in all three cases using Schaefer 200 parcellation. Similarly high agreement was obtained using other outcome measures (e.g., Schaefer 200, matrix correlation: 0.990–0.997; I2C2: 0.982-0.990; discriminability: 1.000). See Supplemental Section S6 for the similarity of intermediate derivatives following replication. It is important to note that these replications did not involve the modification of the original pipelines, so high IPA after replication indicates a faithful replication within C-PAC.

**Figure 2.**
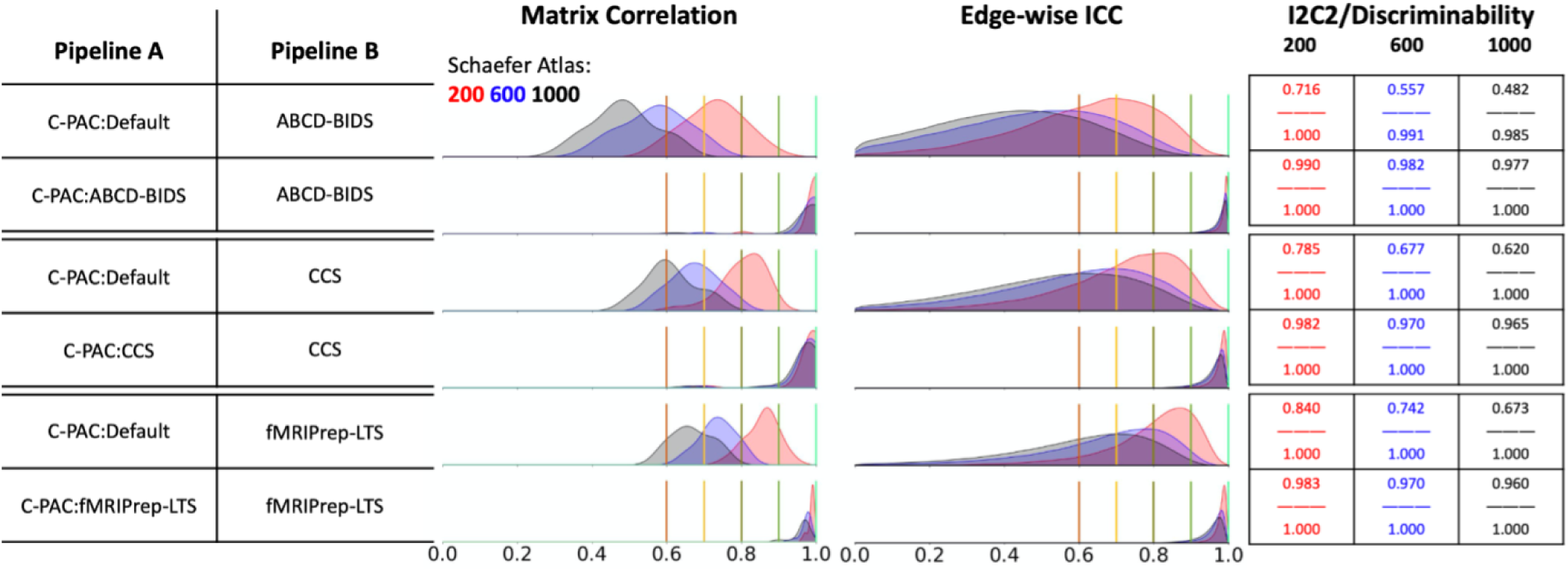
Minimal preprocessing comparisons of C-PAC harmonized pipelines. Each pair of rows shows 1) the agreement between the C-PAC default pipeline and the harmonization target, and 2) the agreement between the C-PAC harmonized pipeline and the harmonization target. The harmonization effort was deemed successful by exceeding an ICC score across pipelines of 0.9, and, in practice, the median ICC score for each harmonized pipeline exceeded 0.98 when using the Schaefer 200 atlas, and 0.96 for the larger parcellations.

### Poor inter-pipeline agreement compromises model replicability in brain-wide association studies

Next, we demonstrated the impact of inter-pipeline agreement on findings in a brain-wide association study (i.e., out of sample classification of biological sex). We examined the comparability of model performance (F1 score) and feature weights obtained in phenotypic prediction tasks when the same datasets were used across four different pipelines (CPAC:ABCD, CPAC:CCS, CPAC:Default, CPAC:fMRIPrep). Given the heavy processing requirements of the pipelines (e.g., Freesurfer in CPAC:CSS, and CPAC:ABCD), we limited our analyses to a sample of 104 datasets (ages: 6.9–16.9 [13.4 ± 2.5]; sex: 56% female) that survived quality control due to head-motion (mean FD ≤ 0.2mm) from a random sample of 300 Healthy Brain Network participants.

We trained a sex prediction model using shared principal components across pipelines to ensure feature comparability, fit on both on uncorrected and age-corrected connectomes to address possible confounds (for more details please refer to the methods section). **Figure 3A** shows that the relative importance of model features (i.e., brain regions of interest) varied substantially across pipelines, despite comparable performance levels across pipelines (as measured by F1-score, **Figure 3B)**, which were also consistent with those reported for sex prediction in the literature^39^. Importantly, the consistency between IPA and feature importance similarity was noted to be extremely strong (Pearson correlation R² = 0.951, p=0.01; **Figure 3C–D**). This demonstrates that while IPA may not be a guiding light in identifying which pipelines may produce the *strongest* associations, it is closely tied to the replicability of results across pipelines.

**Figure 3.**
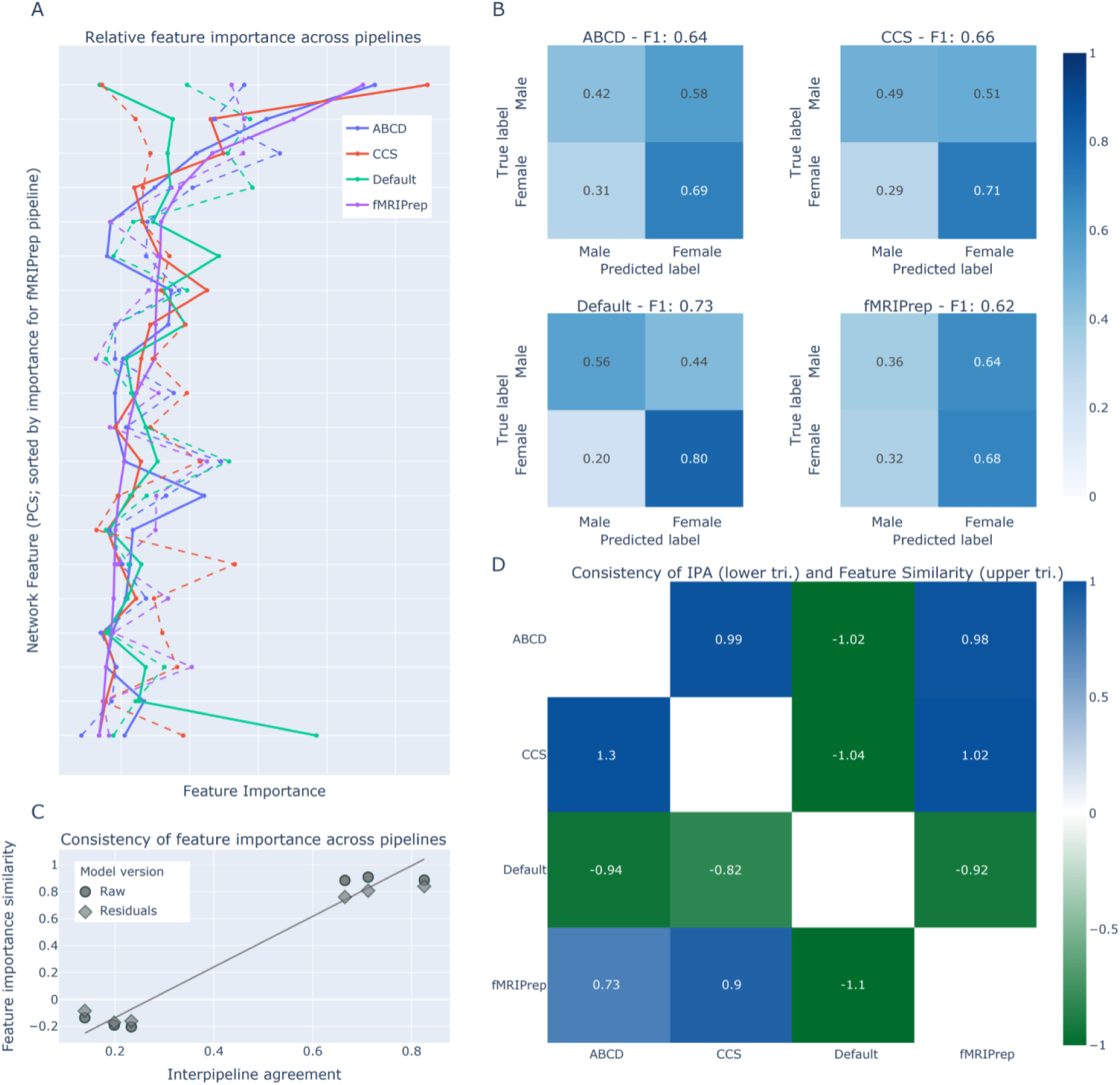
Impact of inter pipeline agreement on result consistency. When using the derived connectivity matrices from each replicated pipeline to predict sex, similar performance was achieved across each processing strategy (B). However, the importance of features in leading to these predictions varied considerably across pipelines **(A),** where solid- and dashed-lines represent the two different modelling approaches. There was a strong relationship (R^2^=0.951, p-0.01) between IPA and the similarity of derived features **(C, D),** demonstrating that variation in pipelines may not significantly impact the strength of discovered relationships, but will regardless lead to variation in the insights made about which regions of the brain, for example, are implicated in such an association.

### Session variation overshadows pipeline differences when scan duration is short

Putting the above findings of pipeline-related variability in the context of known sources of variability is essential when adding these variations to the evolving conceptual models of experimental variation. In this regard, scan duration has emerged as one of the important determinants of test-retest reliability in the literature^40^. In **Figure 4A** we show that the reliability both within and across pipelines was markedly lower for test-retest data than when evaluated with identical data, as would be expected (Kolmogorov-Smirnov test p_corrected_ < 0.001 for all pairwise comparisons). As higher quantities of data per subject were used (i.e., 10 minutes vs 50 minutes), test-retest reliability across sessions dramatically increased both within and across pipelines, from a median edge-wise ICC of 0.227 to 0.611 in the intra-pipeline setting and from 0.152 to 0.428 in the inter-pipeline setting (p_corrected_ < 0.001). In contrast, IPA did not change significantly as the scan duration increased when considering identical data processed by two distinct pipelines (p_corrected_ > 0.1) — this makes sense, as the test-retest reliability for duplicate data is perfect. Taken together, these findings highlight the reality that as the test-retest reliability approaches optimal levels for laboratory measurement, pipeline implementation differences will impose an inherent upper bound on the agreement of preprocessed data. These findings also underscore that 10 minutes of data, which has been common in the field until at least recent years, are insufficient for producing results that are reliable enough to reveal substantive pipeline-related variation.

**Figure 4.**
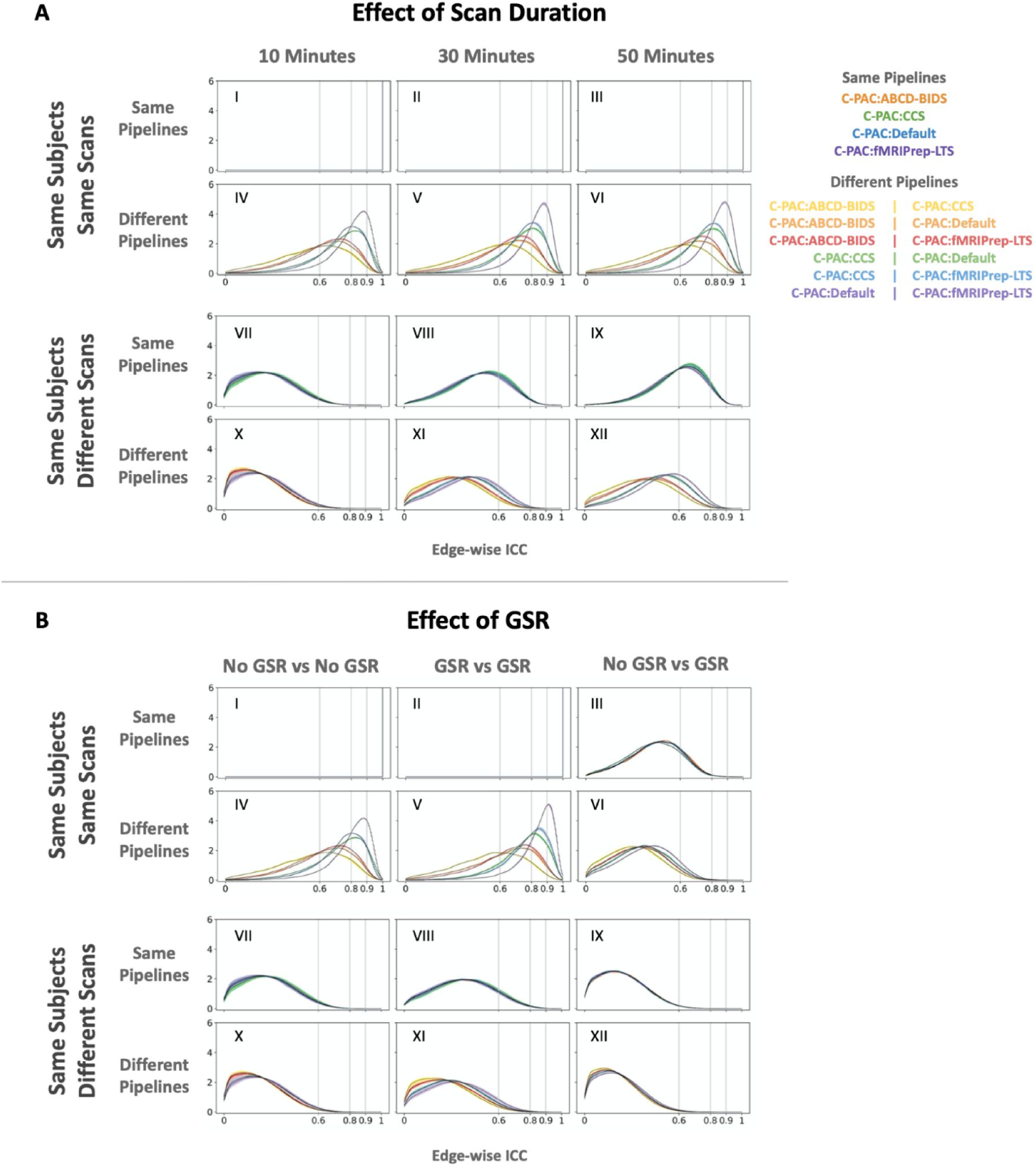
Impact of scan duration (A) and global signal regression (B) on minimal preprocessing results from C-PAC harmonized pipelines. Using both identical (I—VI) and test-retest (VII—XII) data, the ICC both within (l-lll, VII—IX) and between (IV—VI, X—XII) pipelines for each subject was computed. In the case of identical data, both scan duration (A) and the status of GSR (B: columns 1,2) — when matched — had no impact on intra- or inter-pipeline agreement. However, when looking at test-retest data, scan duration was shown to have a significant effect on both intra- and inter-pipeline agreement (A: VII—XII). While the status of GSR did not significantly influence intra- or inter-pipeline agreement, a mismatch in this setting (i.e., only one pipeline using GSR) was found to be highly impactful both when using identical and test-retest data (B: column 3).

### Mismatch in global signal regression is more impactful than minimal processing pipeline differences

Next, we focused on a sometimes-controversial preprocessing step that has varied considerably among resting fMRI studies — global signal regression (GSR)^30^. Specifically, we examined how varying GSR settings affects intra- and inter-pipeline agreement. As shown in **Figure 4B**, when using the same exact 10 minute session data, minimal processing pipeline, and GSR status (i.e., either both "on" or both "off"), perfect agreement was observed; however, median ICC decreased from 1 to notably below the previously mentioned 0.9 threshold when comparing across pipelines — consistent with our findings reported above. A mismatch in GSR (i.e., one pipeline with GSR and the other without) was highly impactful. First, when data and pipelines were matched, a mismatch in GSR resulted in dramatic reductions in IPA (see **Figure 4B**, Panel III), with median ICCs falling below 0.6 (p_corrected_ < 0.0001). In contrast, when using test-retest data (see **Figure 4B**, Panel IX), GSR mismatch effects were more subtle, though still detectable (p_corrected_ < 0.001), with session-related variation being the dominant factor. Relevant to the suggestions of prior work^30,41,42^, IPA was marginally greater when comparing pipelines that both used GSR than pipelines that did not — reaching significance for 3 out of the 6 inter-pipeline comparisons (Mann-Whitney U test p_uncorrected_ = 0.025 – 0.24).

### Spatial normalization workflows typically serve as the biggest source of inter-pipeline variation across minimal processing pipelines

Replicated implementations of the different pipelines in the C-PAC framework afforded us the opportunity to examine which step(s) led to the most variability across pipelines. For each pipeline (C-PAC:Default and C-PAC harmonized versions of ABCD-BIDS, CCS, and fMRIPrep-LTS), we generated a set of pipelines that were each systematically varied by one key processing step across four categories: anatomical mask generation, anatomical spatial normalization, functional mask generation and functional co-registration. Minimal effects were observed when varying denoising components (e.g., non-local means filtering^43^, N4 bias field correction^44^), so this step was merged with mask generation and registration in our evaluation. Each perturbation moved pipelines in the direction of one of the other core pipelines by one component, ultimately producing a space of 48 configurations. As can be seen in **Figure 5**, the specific steps that impact the IPA vary as a function of the specific pairing of pipelines being examined and the interaction of these components.

**Figure 5.**
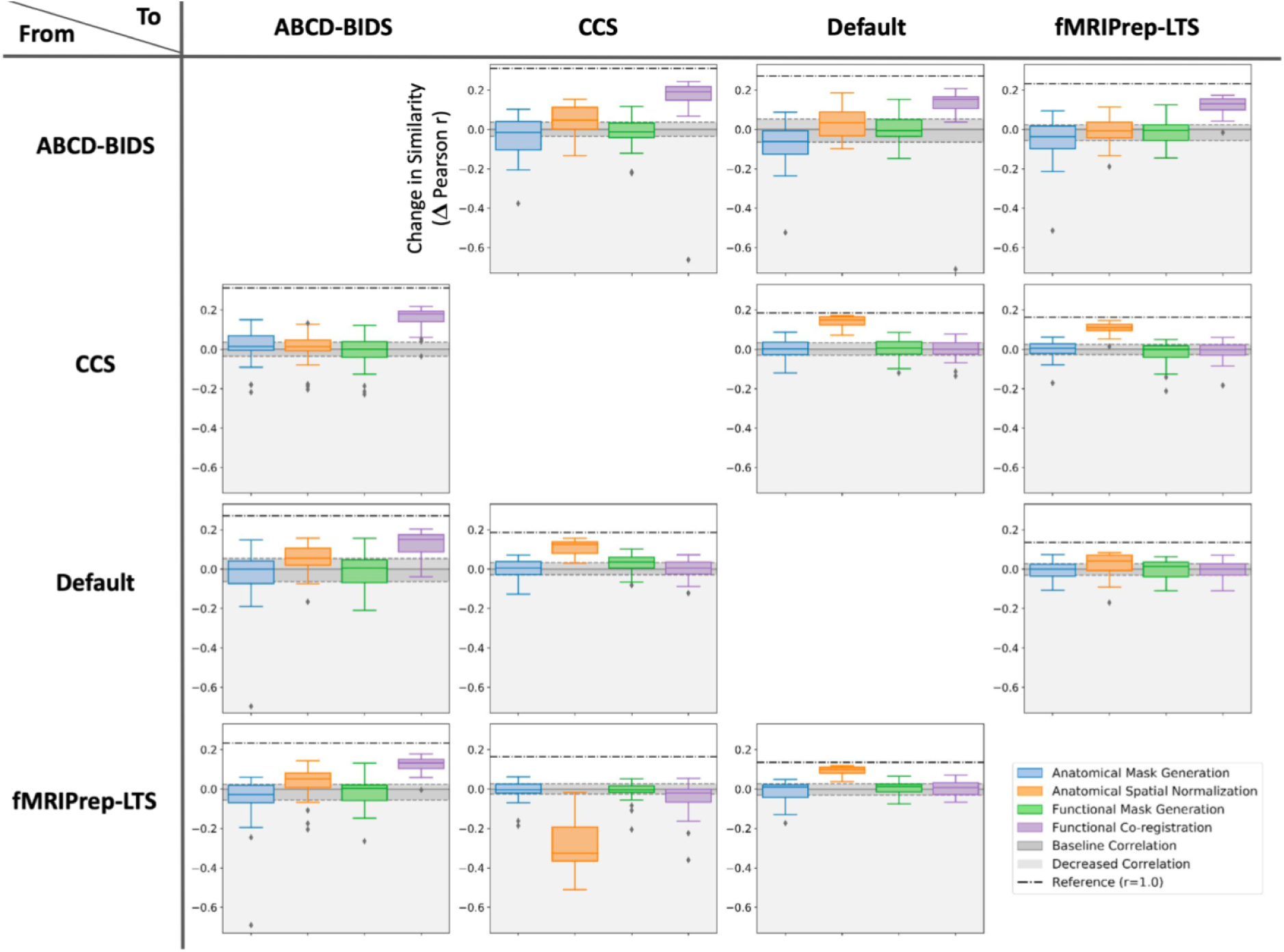
Pairwise identification of sources of variation across harmonized pipelines. Similarity is shown as the difference in Pearson correlation between functional connectivity matrices between the original (harmonized) and perturbed pipelines. Each plot shows the similarity across tools when modifying a single component in the "From" pipeline (rows) to match that in the "To" pipeline (columns). For each pair of pipelines, the zero-line indicates the baseline correlation between the pair before any modifications, and the dashed line indicates the difference between the baseline correlation and a perfect (reference) correlation, i.e., Pearson r = 1.0. Notably, no single step perfectly resolved differences across pipelines, and, in some cases, increasing the component-wise similarity had a negative effect on the agreement of results.

Interestingly, each processing step led to impactful differences in at least one pair of pipelines. However, anatomical spatial normalization and functional co-registration emerged as being among the most consistently impactful (Kolmogorov-Smirnov test p_corrected_ < 0.001 for both spatial normalization steps; p_corrected_ > 0.5 for both mask generation steps). This finding is consistent with the results presented in Figure 1, which illustrate how greater variability is observed across tools at higher-resolutions, namely, that finer-grained parcellations are more acutely affected by registration differences. Importantly, no single step was able to bridge the gap across two pipelines entirely. This is likely a reflection of the complexity of interactions among steps in the pipelines, as well as the possibility that one or more steps other than those examined in this analysis may also be driving findings, such as differences in how spatial transformations are applied to the functional time series (e.g. single-step versus concurrent). For the three pipelines that were closest to one another from the outset (CCS, C-PAC:Default, fMRIPrep-LTS), the anatomical spatial normalization workflow was the biggest determinant of variation. A subtle but important detail of this analysis was that matching of normalization workflows, for example, was not just a matter of matching the registration algorithm, but parameters such as the template resolution, template version, and denoising workflows as well. In addition, matching the functional co-registration step in the ABCD-BIDS pipeline to other pipelines significantly improved the IPA (see Supplemental Section S4). It demonstrates that the BBR option is the biggest source of variation between ABCD-BIDS and other pipelines. Evaluations of the impact of motion correction are shown in Supplemental Sections S7. Of note, increasing the component-wise similarity doesn’t improve the agreement of results in some cases. For example, the correlation decreased when changing the anatomical spatial normalization tool in fMRIPrep-LTS from ANTs to FSL used in CCS. This finding illustrates the complexity of the processing pipelines and shows how their interactions influence pipeline performance.

### Selection of template version and write-out resolution has considerable impact, even within packages

Throughout the pipeline comparison and replication process, we considered various parameter decisions made by users that are not commonly discussed or changed from a pipeline’s default behavior. Of particular note were differences in the specific version of the nearly ubiquitous MNI template and the final write-out resolution of 4D time series, both of which are rarely reported in the literature^45^. In this regard, fMRIPrep-LTS was most disparate, as the default behavior is to write-out using the native image resolution of the fMRI time series (as opposed to 2 or 3 mm-isotropic), and to use the more sharply defined MNI152NLin2009cAsym^46^ template (here referred to as: MNI2009) for reference (as opposed to the MNI152NLin2006Asym^47^ used by most others; here referred to as: MNI2006). To quantify the effects of these seemingly innocuous decisions, even within a pipeline package, we systematically varied them within fMRIPrep-LTS (replicated in the CPAC:fMRIPrep pipeline). As demonstrated in **Figure 6**, while the MNI152Lin^48^ (here referred to as: MNI2001) and 2006 versions of the MNI template generally lead to consistent results, especially when matching output resolution, the 2009 template was markedly distinct. The best case when comparing results generated with the 2009 template and native write-out resolution (default fMRIPrep configuration) and another template were achieved with either the 2001 or 2006 template at a 2 mm isotropic write-out resolution. However, these combinations achieved only a median ICC of 0.89 using the Schaefer 200 parcellation, while the best comparison between the 2001 and 2006 templates maintained an ICC of 1.00. From one perspective, these findings are not surprising given the widespread use of nonlinear registration algorithms, which increase template dependencies. These results nonetheless underscore the impact that even seemingly minor differences in the parameter choices can have substantial implications for intra-pipeline agreement, and would be expected to cascade when considering IPA. One possible limitation of this analysis could be in the quality of transformation of the originally surface-based Schaefer parcellations to the 2009 template^49^; to combat this, we evaluated the correlation of voxelwise time series using each possible pairing of templates (Supplemental Section S8).

**Figure 6.**
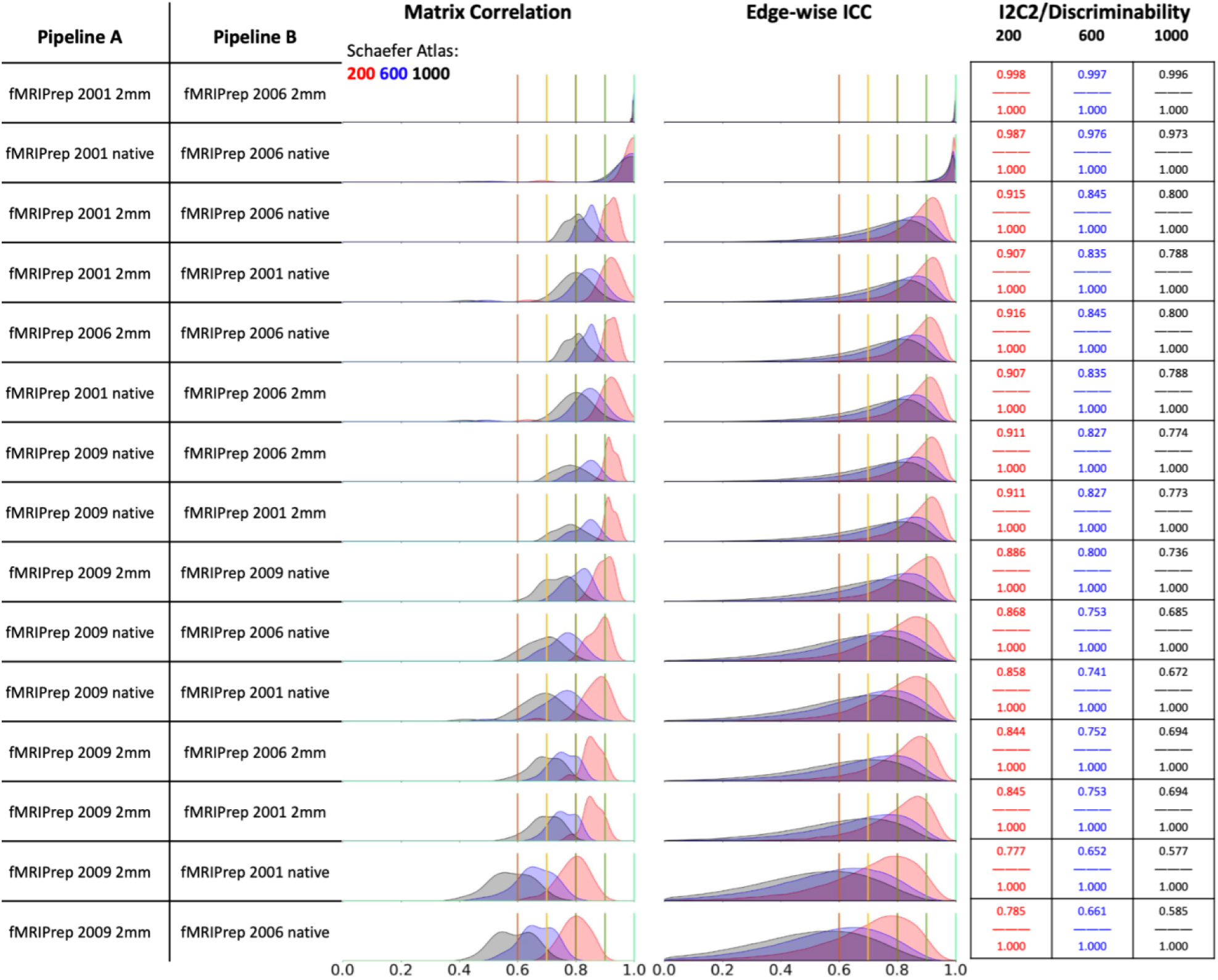
Impact of MNI152 template version and write-out resolution on functional connectomes. Sorted by median individual-level matrix Pearson correlation, the intra-pipeline agreement for fMRIPrep-LTS is shown as both the write-out resolution and version of the MNI152 template are varied. The three most widely used versions of the MNI152 template were used, as well as both 2mm and native-resolution write-out resolution. Agreement is highest across the 2001 (MNI152Lin42) and 2006 {MNI152NLin2006Asym43) versions of the MNI152 template, first when write-out resolution is matched, followed by when it is unmatched across configurations. Below all combinations of 2001 and 2006 templates and write-outs, the default configuration of fMRIPrep-LTS using the 2009 version {MNI152NLİn2009cAsym44) and native write-out resolution is the most highly similar to the others, achieving a median ICC of 0.89 on the Schaefer 200 parcellation. See Supplemental Section S5 for a similar evaluation on voxel-wise time series.

## Discussion

The present work highlights marked variation in individual-level estimates of functional connectivity based on outputs from widely-used functional MRI preprocessing pipelines. Consistent with prior work^20,22,24^, our comparison of minimal preprocessing outputs from five distinct fMRI preprocessing pipelines demonstrated suboptimal Inter-Pipeline Agreement (IPA) for individual differences in functional connectivity, even when identical data was used. Although concerning in the long-term, our analyses using test-retest data suggested that variation arising from insufficient data volume (i.e., short scan durations), which has dominated the literature until recent years, is a more impactful factor than pipeline-related variation at present. Similarly, the present work noted that differences in denoising strategies among studies, such as in whether they include global signal regression, a once highly contested and context-relevant step that comes after minimal preprocessing, can exacerbate pipeline related variation; this finding again emphasizes the need for care in synthesizing the emerging literature focused on individual differences. Perhaps more surprising was our finding that even the most commonly understated of decisions, including the version of the widely used MNI standard space, and write-out resolution, were found to have the potential to pose real limits to intra-pipeline agreement, and more acutely IPA. No one minimal preprocessing component was found to be the dominant source of variation across all pairs of pipelines; instead, the specific steps that most contributed to differences were found to vary depending on which pipelines were being compared. Despite such a broad space of sources for divergence, we demonstrated that variation across pipelines can be overcome through careful replication.

The variations in results arising from pipeline implementation differences in the present work represent an underappreciated bound on the reliability or consistency of results across studies. The impact of implementation differences on IPA were prominent in our analyses, regardless of which pipelines were being compared. Not surprisingly, DPARSF, which is the most distinct with respect to the algorithms and codebase used in its implementation (i.e., SPM/MATLAB-based components), consistently had the lowest IPA across all tested comparisons. Importantly, we show that due to data quality issues, most studies have not yet been limited by the bound imposed by low IPA. Through these experiments we have attributed variability to tool selections. It is likely that imperturbable components (e.g., optimization engine) may play a significant role in differences, but as these components cannot be changed by users directly, they are considered intrinsic to the tools themselves for the purposes of this study. In fact, compromises in measurement reliability, such as the undersampling associated with traditional (short) scan durations, can go so far as to mask implementation differences entirely. Our results demonstrate how pipeline implementation differences will become the next hurdle towards generating findings which can be reproduced across studies as the measurement reliabilities for data collection are optimized — whether through increased scan duration or improved data quality.

The present work also provides a reminder of the variation in findings that arise from methodological variability even within a package. Specifically, we found that the decision of whether or not to include global signal regression was a major source of *intra*-pipeline variation. This is a particularly poignant example, as inclusion of this step has varied across labs and over time as the field has worked towards a consensus on when the method may be more (e.g., correcting for head motion^42,50^ and accounting for variations in arousal), or less useful (e.g., when studying arousal^51^ or other temporal dynamics^52^). While some scientific contexts, such as those mentioned above, may lead to unambiguous decisions regarding the use of GSR, these decisions are much less clear in the context of data dissemination. In these cases, the inclusion or exclusion of GSR may vary based on the preferences of processing teams, and remains an impactful source of variation. The present work also drew attention to the impact that differences in seemingly minimal decisions, such as write-out resolution and template version, can introduce across independent analysts — even when using identical data and an otherwise consistent software package. While the default configuration for the fMRIPrep-LTS package adopts the newer MNI2009 asymmetric^46^ template, the majority of the field uses the 2006 asymmetric^47^ or 2001^48^ templates, which are more similar to one another. Similarly, fMRIPrep-LTS uses a native write-out resolution, while most other tools use 2 or 3 mm. The write-out resolution and template versions were found to interact and establish tiers of agreement across results generated within fMRIPrep, importantly demonstrating that the effect of these factors is non-uniform. For example, matching either template version or write-out resolution alone may not lead to the highest similarity of results. The fact that using higher resolution atlas results in lower IPA also shows the sensitivity of atlas-based connectivity analysis. Improving registration consistency within regions of interest can be a vital step towards higher reliability, but is of course downstream of improving registration consistency as a whole for the purpose of whole-brain analysis.

It is important to note that greater agreement across pipelines does not necessarily imply greater validity or quality of the results. For example, the DPARSF pipeline uses SPM’s DARTEL registration tool, which is known to be a high quality and reliable tool for spatial normalization^17^, but it has low agreement with other pipelines. While the present work primarily focused on measures of reliability for its evaluation, a critical prerequisite for usage of tools, future work would benefit from using validity as a primary target (e.g., predictive/explanatory power). This is a logical order of examination for these two constructs, as reliability, either across measurements or methodological choices, places an upper bound on validity or utility.

Perhaps the most poignant finding of this study was that low IPA also compromises the insights derived from data when used in downstream analyses. Our BWAS experiment showed that there was a strong relationship between IPA and feature importance when constructing a model for an association/prediction task — providing an empirical reminder that reliability bounds validity. This means that any interpretation applied to which features (i.e., functional connections) led to the model performance would be unlikely to replicate across pipelines, when IPA is low. As brain imaging seeks to advance our basic understanding of the brain, this result is crucial as it reminds us that each individual pipeline allows us to test a specific, but not unique, version of a hypothesis relating brain organization to a question of interest. It is therefore important to not merely adapt a single pipeline when performing such work, but to adopt several.

A potential limitation of the present work could be that it was largely carried out on a dataset acquired with traditional MRI protocols of standard data quality, popular at the time of acquisition, rather than increasingly adopted but historically inaccessible practices (e.g., inclusion of distortion field maps). Two key factors drove this decision. First, the limited availability of high characterization, test-retest datasets that employ more modern acquisition methods. The larger bulk of sufficiently-powered test-retest data available to date are either single band EPI data or do not exceed 60 minutes total per subject. While the Midnight Scan Club^53^ collection has higher quality data for each individual, it is limited to a cohort of only 10 participants. Second, the data employed are representative of, or exceed, the quality of the data employed in the majority of fMRI datasets available to most researchers — particularly those in clinical populations (e.g., ABIDE^54^). Our supplementary analyses using both a low- and high-motion cohort of participants from the Healthy Brain Network (HBN) dataset^37^, which includes imaging data obtained using state-of-the-art functional MRI (i.e., multiband) and anatomical sequences from the NIH ABCD Study^55^, showed similar IPA to those obtained with the HNU dataset, though slightly improved (Supplemental Section S5). This tempers hopes that improving data quality alone will improve agreement between differing processing pipelines, or analytic tools more broadly. Another limit of this work is that the sole denoising strategy evaluated was global signal regression. Our motivation for exploring GSR alone was not to imply that it is the solely significant denoising step in processing pipelines; rather, we aimed to provide a known and widely recognized benchmark for variation across tools that could be used to situate the observed variation across processing pipelines. Other denoising approaches such as white matter and CSF mask regression, respiratory and cardiac noise removal and ICA-AROMA can also potentially impact preprocessing results and merit dedicated examination in future work.

The data and pipelines tested here represent a snapshot of those that have been or will be used in the field of functional neuroimaging. However, they also illustrate challenges to reproducibility that are likely pervasive across various modalities of brain imaging research — or computational science more broadly, — and will continue to be faced as fields move forward. We show that modernization of techniques alone (e.g., improvement of data quality) will not overcome challenges related to IPA and can actually amplify them (e.g., improvements in MNI template quality). In addition, our findings motivate a number of considerations for how to improve reproducibility. Arguably, the most easily actionable, is for publications to include rich and detailed specifications of all data processing software (e.g., tool versions, parameters, templates, in-house code) to promote reproducibility, ideally, with all code being made available through a public repository (e.g., GitHub, Zenodo). Beyond this, there is a need for the field to increase its focus on testing of tools, and benchmarking of novel pipelines against one or more reference pipelines (e.g., fMRIPrep-LTS, HCP Pipelines). Adopting evaluation standards consistent with computer science and industry will not only increase the transparency of tools and results, but provide greater context for their relationships with one another. Until clear benchmarks and bridges between tools can be established, one recommendation for authors is to repeat their analyses with an auxiliary pipeline, or by perturbing template version, write-out resolution, or other specific analytic decisions (specified prior to initiation of a research project), and reporting potential dependencies of results on the selected pipeline. Lack of replication would not necessarily undermine the value of results obtained with any one pipeline, but rather draw attention to potential dependencies that can limit reproducibility if not taken into consideration. The C-PAC framework makes the process of using multiple pipelines relatively easy for scientists. Beyond using multiple pipelines, strategies for consolidating results across pipelines should be identified. Depending on the analytic goals, this could involve the aggregation of results (e.g., bagging) to generate composite findings, or the ensembling of results to improve prediction^56,57^. This has been recently demonstrated in brain imaging and numerical uncertainty^58–60^.

Focused on the optimization of test-retest reliability within-pipelines over the past decade, the functional neuroimaging field now needs to take on a new major challenge — Inter-pipeline agreement. The present work draws attention to the substantial impact that variations in the most basic processing steps can introduce into imaging results. The challenges and solutions provided in the present work are not specific to neuroimaging, but instead, representative of the process that the broader field of neurosciences will need to go through to become a reproducible science.

### Online Methods

#### Dataset

Analyses in the present study were carried out using the Hangzhou Normal University (HNU) test–retest dataset made publicly available via the Consortium for Reliability and Reproducibility^31^ (CoRR). The dataset consists of 300 R-fMRI scans, collected from 30 healthy participants (15 males, age = 24 ± 2.41 years) who were each scanned every three days for a month (10 sessions per individual). Data were acquired at the Center for Cognition and Brain Disorders at Hangzhou Normal University using a GE MR750 3 Tesla scanner (GE Medical Systems, Waukesha, WI). Each 10-min R-fMRI scan was acquired using a T2*-weighted echo-planar imaging sequence optimized for blood oxygenation level dependent (BOLD) contrast (EPI, TR = 2000 ms, TE = 30 ms, flip angle = 90°, acquisition matrix = 64 × 64, field of view = 220 × 220 mm^2^, in-plane resolution = 3.4 mm × 3.4 mm, 43 axial 3.4-mm thick slices). A high-resolution structural image was also acquired at each scanning session using a T1-weighted fast spoiled gradient echo sequence (FSPGR, TE = 3.1 ms, TR = 8.1 ms, TI = 450 ms, flip angle = 8°, field of view = 220 × 220 mm, resolution = 1 mm × 1 mm × 1 mm, 176 sagittal slices). Foam padding was used to minimize head motion. Participants were instructed to relax during the scan, remain still with eyes open, fixate on a displayed crosshair symbol, stay awake, and not think about anything in particular. After the scans, all participants were interviewed to confirm that none of them had fallen asleep. Data were acquired with informed consent and in accordance with ethical committee review. One subject sub-0025430 was excluded in all analyses because of its inconsistent preprocessed results across all pipelines. For supplemental analysis on a higher quality dataset using the state of the art sequences, we used 29 high-motion and 29 low-motion subjects of the Healthy Brain Network (HBN) dataset^37^; for more information, please see the referenced publication. Importantly, as noted in a recent work^61^, the sample size employed in the present work is more than sufficient to detect moderate–excellent ICC’s (e.g., 0.5 and higher) if using two scans per subject, and well powered for notably smaller effects when using the larger number of scans per individual.

#### Assessment of inter-pipeline agreement

Five pipelines were used to measure IPA - ABCD-BIDS v2.0.0, CCS version in May 2021, C-PAC:Default v1.8.1, DPARSF v4.5_190725, fMRIPrep-LTS v20.2.1. We pursued a multifaceted assessment strategy to evaluate test-retest reliability, including: 1) Individual-level matrix Pearson correlation of functional connectivity matrices across pipelines, 2) the edge-wise intra-class class correlation coefficient (ICC), 3) the image intraclass correlation coefficient^32^ (I2C2, connection-wise index of reliability), and 4) discriminability^33^ (matrix-level index of reliability). For each of these measures, we evaluated multiple scales of spatial resolution (200, 600, 1000 Schaeffer parcellation units) to explore the relationship of results with the number of parcels. The Schaeffer atlas was resampled to the output space using FSL FLIRT accordingly and then parcels were extracted using AFNI 3dROIstats. IPA (matrix correlation, ICC, I2C2 and discriminability) was evaluated for both a) across different sessions and b) across different pipeline configurations (using identical data).

#### Replication process

First, we surveyed the ABCD-BIDS, CCS, C-PAC and fMRIPrep-LTS pipelines for differences in which steps and libraries were included. We identified preprocessing components (e.g., motion correction as implemented by FSL^62^ MCFLIRT^63^) which were not found in C-PAC and added them to the codebase. Key differences identified in this process are depicted in **Table 1**. For all replication exercises, the C-PAC default pipeline was used as a base which was modified iteratively. While the ultimate goal of the replication process was to achieve connectivity matrices with a correlation of 0.9 or higher across all measures, we examined a range of intermediates to facilitate the implementation and debugging process (see Supplemental Section S6 for a list of intermediates, and **Figure 1** for sample comparison indicator boards generated to guide process).

#### Anatomical preprocessing differences and replication

The major components of the C-PAC default anatomical workflow is as follows: 1) brain extraction, via AFNI^64^ 3dSkullStrip; 2) tissue segmentation, via FSL FAST^65^; and linear and non-linear spatial normalization, via ANTs^66^. The ABCD-BIDS pipeline applies extensive preprocessing prior to brain extraction, including: non-local means filtering^43^, N4 bias field correction^44^, Anterior Commissure - Posterior Commissure (ACPC) alignment, FNIRT-based brain extraction, and FAST bias field correction. Following these steps, FreeSurfer^67^ is used for brain extraction and segmentation masks are refined prior to image alignment using ANTs. The CCS pipeline also applies non-local means filtering on raw anatomical images and uses FreeSurfer to generate the brain and tissue segmentation masks, followed by linear and non-linear alignment using FSL and skull-stripped images. Note that CCS is the only pipeline using FSL for image registration. The fMRIPrep-LTS pipeline applies N4 bias field correction, followed by ANTs for brain extraction, a custom thresholding and erosion algorithm to generate segmentation masks, and ANTs for image registration. In the case of fMRIPrep-LTS, ANTs registration is performed using skull-stripped images, unlike C-PAC default and ABCD-BIDS which use whole-head images. Note that we opted to use the volume-based fMRIPrep-LTS workflows rather than surface, to increase the similarity with other pipelines, which are primarily focused on volume-space analysis.

#### Functional preprocessing differences and replication

The major components of the C-PAC default functional preprocessing workflow is as follows: 1) slice timing correction, via AFNI 3dTshift; 2) motion correction, via AFNI 3dvolreg; 3) mask generation, via AFNI 3dAutomask; 4) co-registration with mean function volume, via FSL FLIRT^63,68^; 5) boundary-based registration, via FSL FLIRT; 6) time series resampling into standard space, with ANTs. The ABCD-BIDS pipeline uniquely does not perform slice timing correction, and is the only pipeline which does not use boundary-based alignment when no distortion map is provided. Further, ABCD-BIDS resamples the anatomical mask to the functional resolution. The CCS pipeline implements despiking with AFNI 3dDespike as the first functional preprocessing step, and the functional mask is generated by further processing of the anatomical brain mask. The fMRIPrep-LTS pipeline also implements despiking with AFNI 3dDespike as the first functional preprocessing step, and uses a hybrid AFNI-FSL brain extraction approach for mask generation. Interestingly, there are two steps in which no two pipelines are identical: mask generation and co-registration. In the case of mask generation, four distinct approaches are used, while for co-registration, four different functional volumes are selected in four pipelines. At the final time series resampling step, both ABCD-BIDS and fMRIPrep-LTS use a similar one-step resampling approach to apply motion correction, co-registration and anatomical to standard-space registration matrices simultaneously, while CCS and C-PAC apply transformations on the functional times series sequentially.

By replicating the key methodological choices from each of the pipelines, we were able to implement ABCD-BIDS, CCS and fMRIPrep-LTS pipelines in C-PAC (referred to as C-PAC:ABCD-BIDS, C-PAC:CCS, C-PAC:fMRIPrep-LTS).

#### Impact of pipeline variation on prediction

We conducted a comparative analysis among distinct data processing pipelines to assess the influence of pipeline variability on a sex prediction task. To limit the processing requirements for this analysis, we limited our analyses to a random sample of 300 Healthy Brain Network participants, using only those that survived quality control due to head-motion (mean FD ≤ 0.2mm). Specifically, we used 104 resting-state scans with minimal head motion from the HBN project and processed them using four pipelines (C-PAC:Default, CPAC:ABCD-BIDS, C-PAC:CCS, C-PAC:fMRIPrep-LTS). After processing the data, we computed the connectivity matrices for each subject, outlining interactions across 200 predefined brain regions from the Schaefer200 atlas. Subsequently, we employed a linear regressor to remove the age-related signal for each edge within each pipeline, resulting in two sets of connectomes per-pipeline: age-corrected and age-uncorrected. Then we built a sex predictor by selecting the 20 most important components for each pipeline. To evaluate the impact of pipeline variations on prediction outcomes, we performed stratified 10-fold cross-validation using a random forest classifier under four distinct conditions: 1) applied a common PCA approach, concatenating and computing principal vector collectively for all pipelines; 2) a similar PCA method but on the residuals obtained from fitting a linear regression model to each edge and age, pipeline-wise; 3) an individual PCA approach, computing principal vector independently within each pipeline; and 4) same as the previous, but involving the PCA on the residuals of linear regression. The results of stages 1 and 2 were used to perform the consistency analysis, while stages 3 and 4 were used to ensure consistency in performance in a "native", unpooled pipeline setting.

#### Impact of scan duration on intra- and inter-pipeline agreement

We repeated our inter-pipeline comparisons to evaluate the role that scan duration plays on the reproducibility of the results across minimal preprocessing configurations. Ten comparisons were made, consisting of random samples of 10-, 30- and 50-minutes of fMRI data per subject generated from the HNU test-retest dataset, in which each scan contains 10-minutes of fMRI data per subject. The impact of scan duration was evaluated with respect to both the cross-scan test-retest reliability and the IPA.

The impact of scan duration was examined under two conditions. First, the exact same data was used in each pipeline (i.e., same subjects, same sessions). This provided a condition of perfect data test-retest reliability, thereby allowing examination of pipeline differences in isolation of any compromises related to the data. In the second condition, we used non-overlapping scan data from the same subjects to each pipeline, allowing us to observe the collective compromises in reliability related to the data and the pipelines. By varying combinations of scan and pipeline comparisons at the same time, we arrived at four distinct categories of comparisons: 1) same scan, same pipeline; 2) same scan, different pipelines; 3) different scans, same pipeline; 4) different scans, different pipelines.

#### Impact of global signal regression on intra- and inter-pipeline agreement

The impact of global signal regression (GSR) was assessed under the same conditions as evaluations of scan duration, above. The impact of GSR was evaluated with respect to both the across-scan and inter-pipeline test-retest reliability. To perform GSR, we used functional time series and functional brain masks in template space, ran AFNI 3dROIstats to get the mean time series, and then ran AFNI 3dTproject with quadratic detrending to get GSR time series for each pipeline. For consistency, we repeated our inter-pipeline GSR evaluations 10 times, each using 10-minutes of fMRI data, in each of three settings: **no** GSR vs **no** GSR, GSR vs GSR, and **no** GSR vs GSR. For statistical testing across settings, the distribution of ICC scores were compared to another. A Mann-Whitney U-test was chosen as it’s a non-parametric test to say if samples from one distribution are likely to be higher than another, i.e. if GSR ICC scores are likely to be higher than non-GSR scores.

#### Impact of template version and write-out resolution

In the course of our work, we noted that even the most highly prescribed pipelines allow users to make decisions regarding template version and write-out resolution. To evaluate the potential impact of these decisions, we examined the impact of each on estimates of functional connectivity generated by the same package using different options. We carried these analyses out in both CPAC:fMRIPrep and the fMRIPrep-LTS pipeline configuration without surface reconstruction (i.e., --fs-no-reconall configuration), using templates from TemplateFlow^45^. We selected three templates: the original (linear) MNI152Lin^48^ (here referred to as: MNI2001), MNI152NLin2006Asym^47^ (here referred to as: MNI2006), and MNI152NLin2009cAsym^46^ (here referred to as: MNI2009). The templates were updated in each of the 2006 and 2009 versions using improved alignment algorithms, leading to an increased detail and quality. The 2009 version is used as the default spatial-standardization reference in fMRIPrep-LTS, while the 2001 and 2006 versions are distributed with FSL and used in other pipelines. We evaluated two different write-out resolutions — a 3.4 × 3.4 × 3.4 mm resolution matching that of the native functional images, which is used in fMRIPrep-LTS, and a 2 × 2 × 2 mm resolution used in ABCD-BIDS. This gave us six different processing tracks in total. We then repeated our inter-pipeline agreement measures using functional connectivity matrices from these six processing tracks. We report the fMRIPrep-LTS findings given the widespread use of the package, though note the CPAC:fMRIPrep-LTS findings were identical.

#### Sources of variation

We utilized the configurable options in C-PAC to evaluate sources of variation among four pipelines (C-PAC:ABCD-BIDS, C-PAC:CCS, C-PAC:Default, C-PAC:fMRIPrep-LTS). We first calculated the matrix correlation of functional connectivity matrices using Schaefer 200 parcellation across every two pipelines as a baseline. Each of four pipelines is used as a source pipeline and the other three pipelines are used as target pipelines. We then varied each of the configurable options in the source pipeline to the target pipeline’s option at four key preprocessing steps (anatomical mask generation, anatomical spatial normalization, functional mask generation, functional co-registration), and observed how the change of configuration at each preprocessing step affects the final minimal preprocessing result. Pearson correlation of functional connectivity matrices from the Schaefer 200 parcellation was used to estimate pipeline differences.

#### Statistical analysis

Unless otherwise stated, when distributions of agreement measures were being compared across settings, either a Kolmogorov–Smirnov test^69^ (KS test), Mann-Whitney U-test^70^ (MWU test), or paired t-test were performed. In the case of comparisons where the objective was to test if two distributions were different from one another, KS-tests were used. In contrast, for the cases where the objective was to evaluate if samples from one distribution were more likely to be of a higher value than another, MWU-tests were performed. In the case where comparisons were being made within a given configuration and across parcellations, paired t-tests were used. In all cases, equivalent tests were corrected for multiple comparisons using the highly conservative Bonferroni correction technique.

#### Code availability

All software created and used in this project is publicly available. The C-PAC pipeline is released under a BSD 3-clause license, and can be found on GitHub at: https://github.com/FCP-INDI/C-PAC/releases/tag/v1.8.2; the ABCD-BIDS pipeline is released under a BSD 3-clause license, and can be found at: https://github.com/DCAN-Labs/abcd-hcp-pipeline/releases/tag/v0.0.3; the CCS pipeline can be found at: https://github.com/zuoxinian/CCS; the fMRIPrep-LTS pipeline is released under Apache License 2.0 and can be found at: https://github.com/nipreps/fmriprep/releases/tag/20.2.1. Templates were all accessed through TemplateFlow^45^. All analysis software, including experiments and figure generation, can be found on GitHub at https://github.com/XinhuiLi/PipelineAgreement as well as on Zenodo at https://zenodo.org/badge/latestdoi/415936717. The preprocessed functional connectivity data can be found on OSF at https://osf.io/kgpu2/.

## Acknowledgements.

This work was supported in part by gifts from Joseph P. Healey, Phyllis Green, and Randolph Cowen to the Child Mind Institute. Additionally, grant awards from the NIH BRAIN Initiative to MPM and RCC (R24 MH11480602), to GK and MPM (RF1MH130859), to RP, OE, MPM and TS (RF1MH121867); and from NIMH to TS and MPM (R01MH120482). OE received support from the SNSF Ambizione project 185872. CGY received support from National Natural Science Foundation of China (grant number: 82122035, 81671774, 81630031).

## Supplemental Sections

**S1: Key conceptual differences between pipeline pairs**

The conceptual differences between pipeline pairs can be classified into three categories: (1) Same approach, same objective; (2) Different approach, same objective; (3) Different approaches, different objectives. We compared all combinations of pipeline pairs across five packages. Conceptually different steps, such as co-registration, also lead to significantly different preprocessed results as shown in Figure 5.

**Table S1.1.**
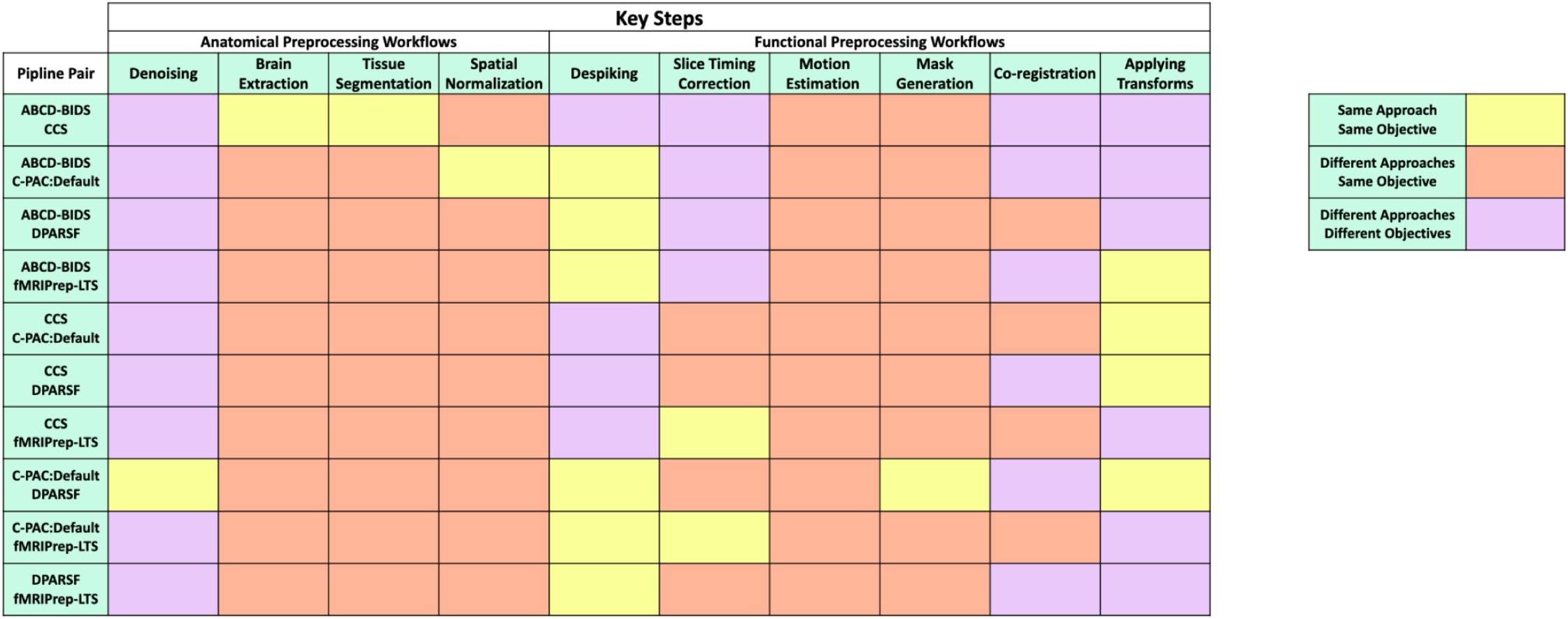
Key conceptual differences between pipeline pairs.

**S2: Inter-pipeline agreement on brain regions**

**Figure S2.1** shows inter-pipeline agreement based on parcellated brain regions. Each plot in the upper triangle is the ICC heatmap, each plot in the lower triangle is the mean ICC (top) and coefficient of variation — the ration between the mean and the variance — of ICC scores for each region (bottom) in each brain parcel region mapped to the parcellated brain (Schaefer 200).

**Figure S2.1.**
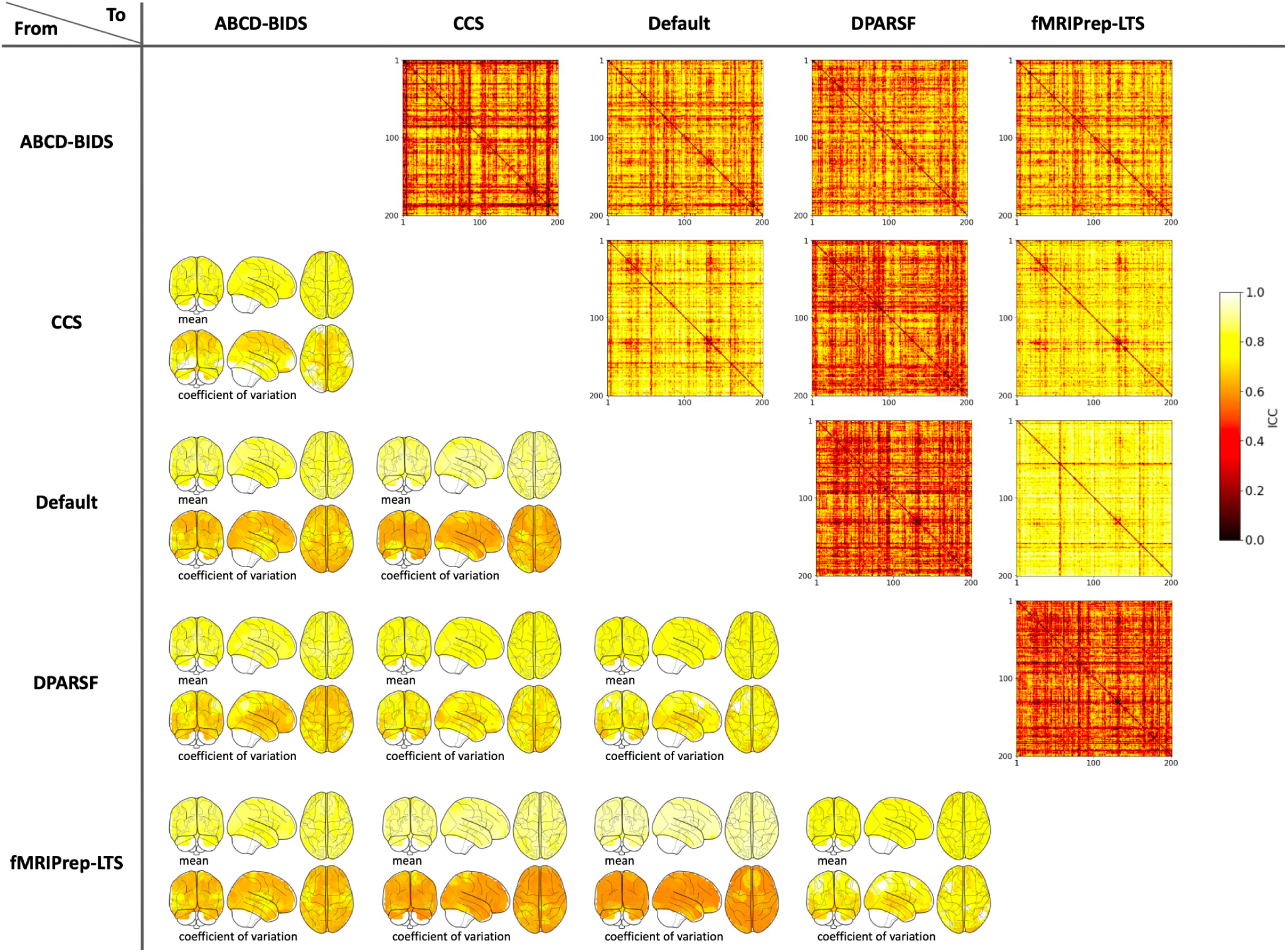
Inter-pipeline agreement on brain regions.

**S3: Impact of pipelines on subject-sorting via graph theory measures**

To evaluate how function connectivity differences from different pipelines are sensitive to graph theory measures, we measured global connectivity and degree centrality with sparsity thresholds as 90%, 95% and 97.5%, respectively, on 29 HNU subjects and 29 HBN low-motion subjects. Though none of the measures shows significant difference in the overall sample distribution across pipelines (**Figure S3.1** and **Figure S3.4**, Kolmogorov-Smirnov test), the pipeline difference can be observed across individuals (**Figure S3.2** and **Figure S3.5**). We further sorted each type of metrics for each pipeline and compared the agreement of the ranked subject lists across pipelines. Specifically, we counted the number of subjects that remain in the same order between pipeline pairs and then divided by the total number of subjects (**Figure S3.3** and **Figure S3.6**). We calculated the average ratio across all pairs for a given measure, and defined this as the average intersection score. The average correlation of subject ranks of global connectivity, along with degree centrality when computed with three distinct sparsity thresholds of 90%, 95% and 97.5%, respectively, are: 0.22, 0.12, −0.03, 0.22 on the HNU dataset, and 0.45, 0.24, 0.13, 0.23 on the HBN dataset, respectively. The average position-difference of above metrics are: 7.26, 8.67, 9.74, 8.20 on the HNU dataset and 5.82, 7.76, 8.55, 7.99 on the HBN dataset, wherein an ideal ranking difference would be 0 and the maximum average difference would be 14.48. The result demonstrated that pipeline-related variation will result in significant differences across individuals on graph theory measures, and subsequent downstream analyses.

**Figure S3.1.**
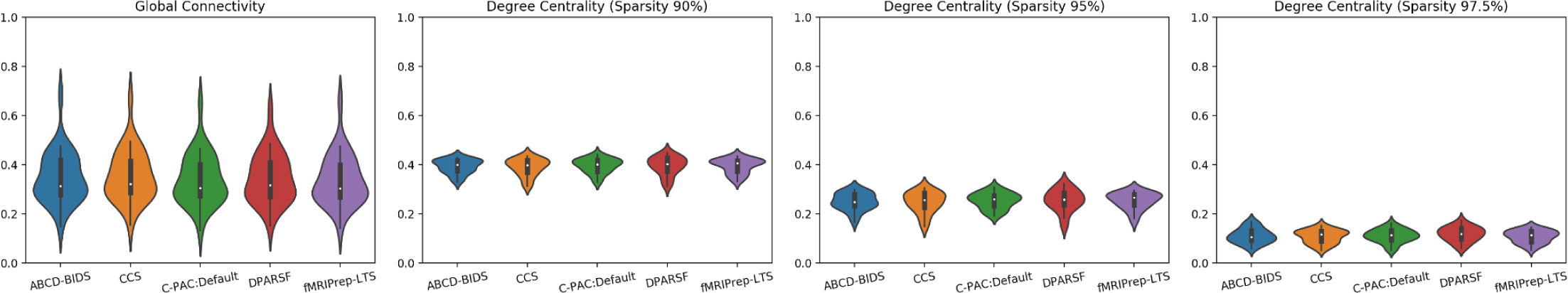
Graph theory measures of pipelines on the HNU dataset.

**Figure S3.2.**
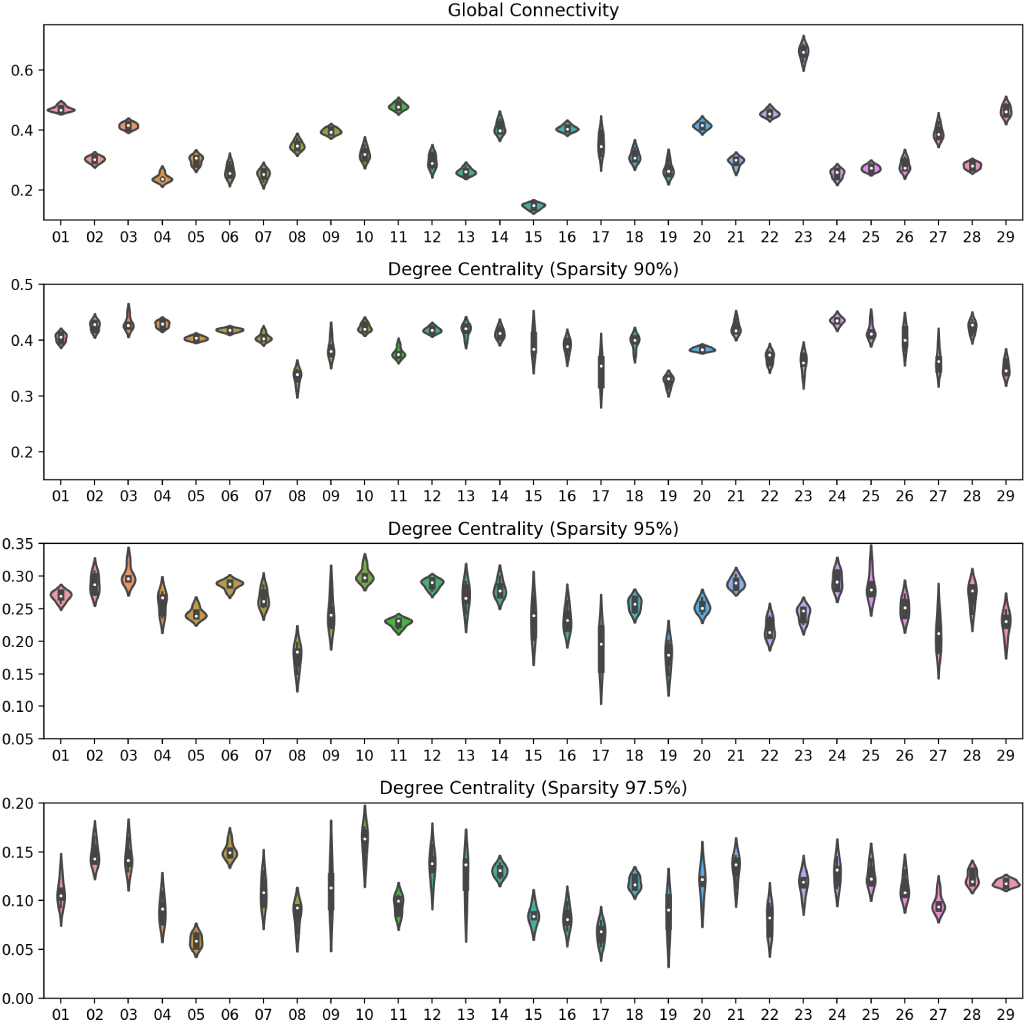
Graph theory measures of individuals on the HNU dataset.

**Figure S3.3.**
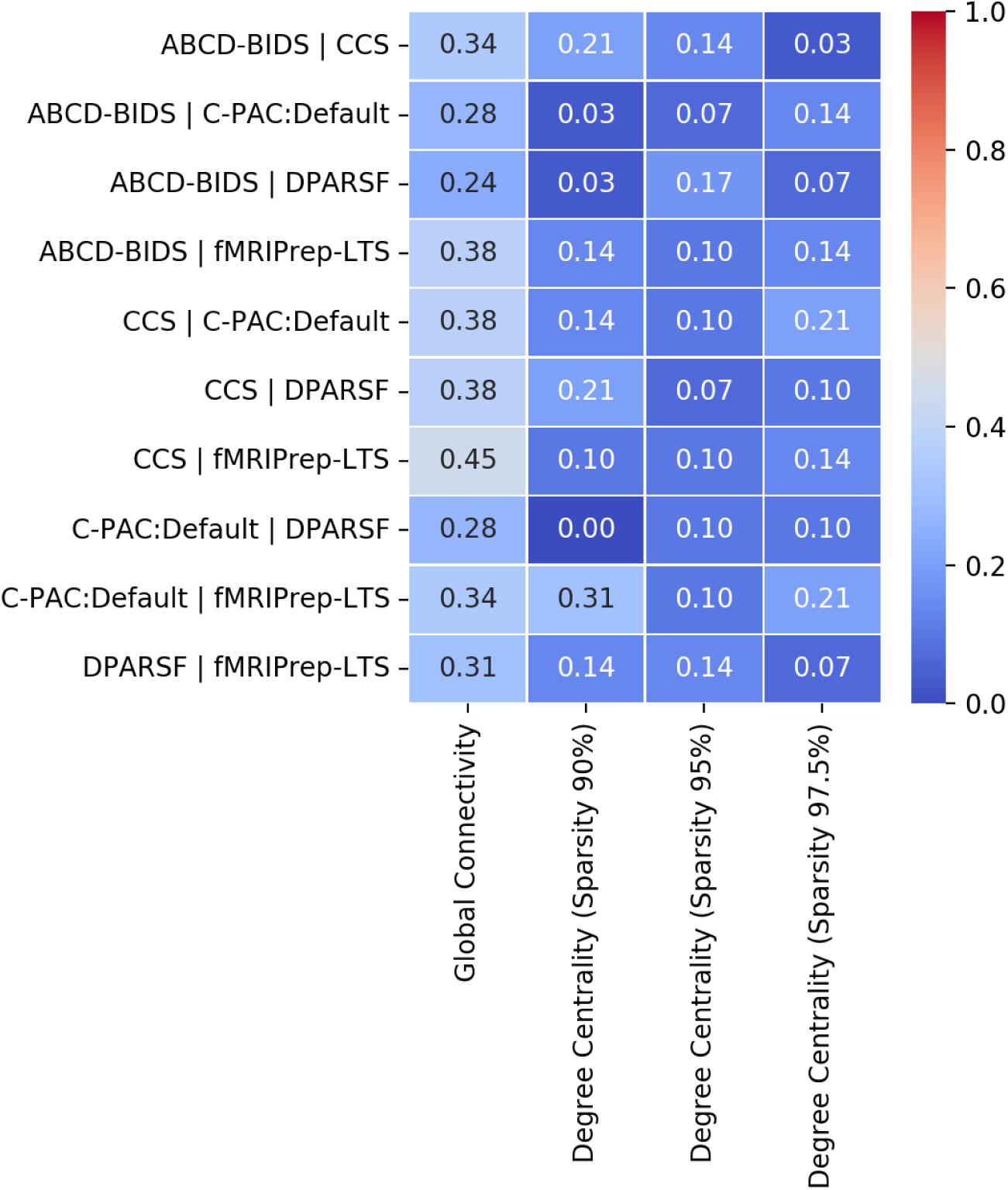
The ratio of intersected subjects between pipeline pairs on the HNU dataset.

**Figure S3.4.**
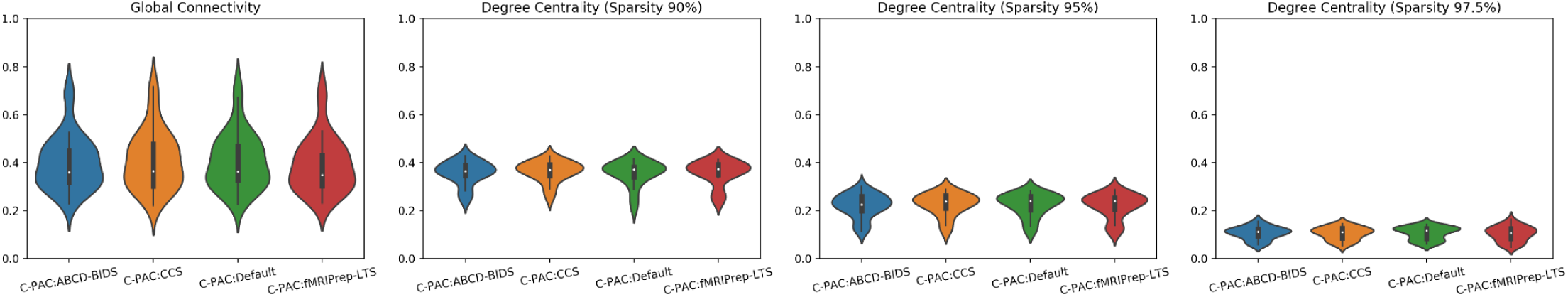
Graph theory measures of pipelines on the HBN dataset.

**Figure S3.5.**
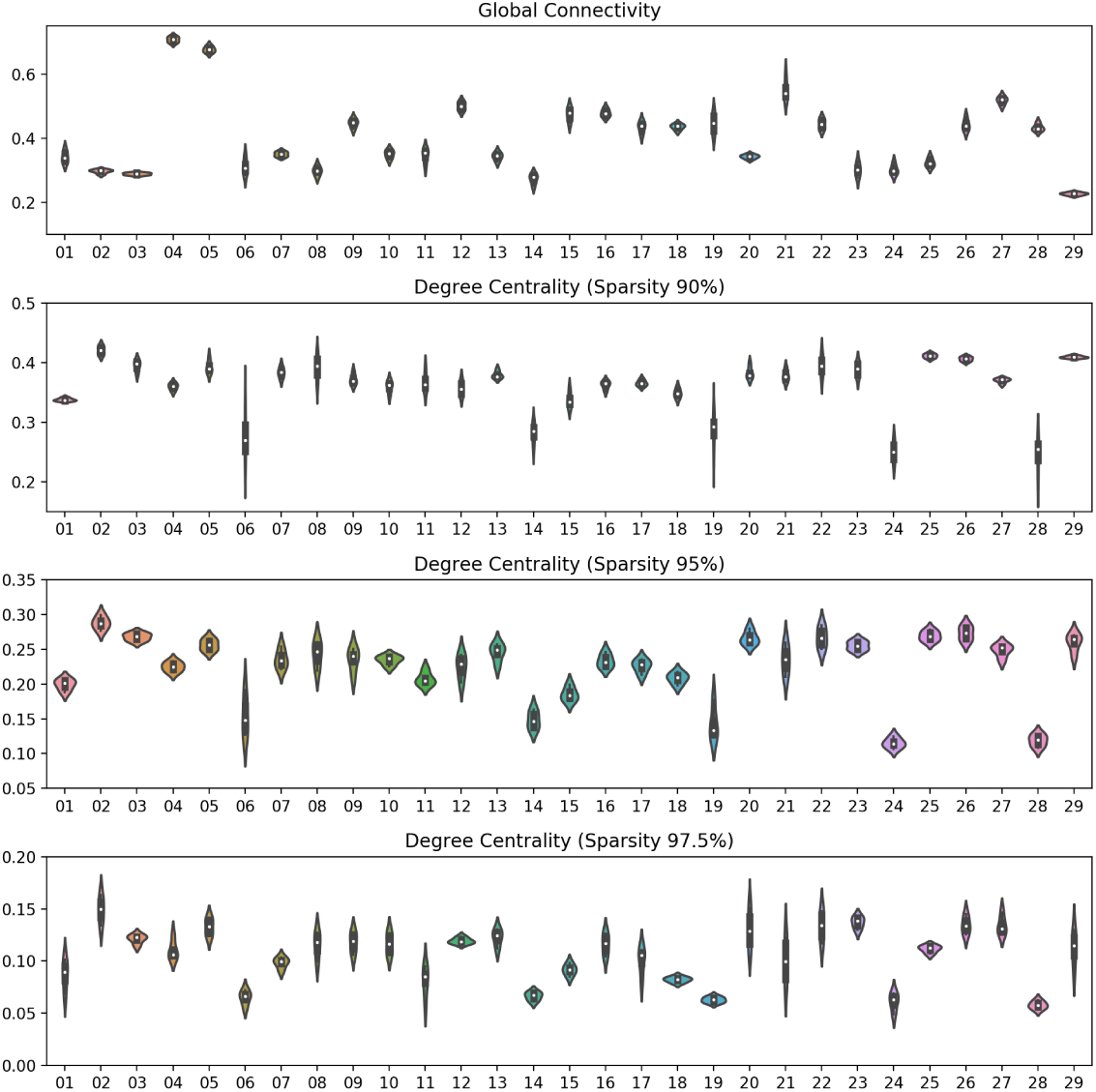
Graph theory measures of individuals on the HBN dataset.

**Figure S3.6.**
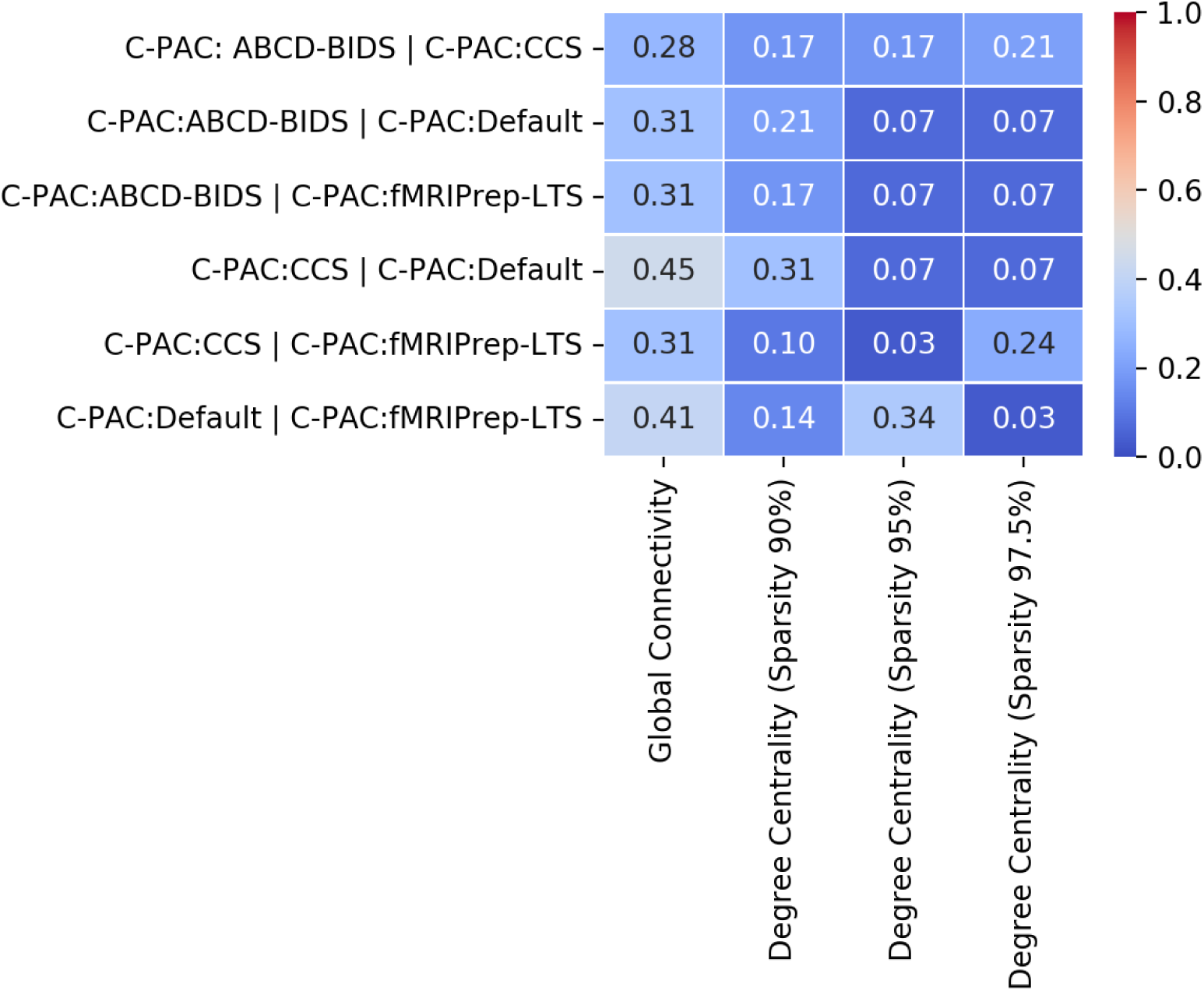
The ratio of intersected subjects between pipeline pairs on the HBN dataset.

**S4: Modifying C-PAC:ABCD-BIDS to use boundary-based registration**

To confirm that the boundary-based registration (BBR) step is the main source of variation in the ABCD-BIDS pipeline, we utilized the configurable option in C-PAC and turned on the BBR step in C-PAC:ABCD-BIDS to generate the C-PAC:ABCD-BIDS BBR pipeline. We then repeated the inter-pipeline reliability measures among five pipelines (C-PAC:ABCD-BIDS BBR, CCS, C-PAC:Default, DPARSF, fMRIPrep-LTS). **Figure S4.1** indicates that the IPA between the C-PAC:ABCD-BIDS BBR pipeline and other pipelines improves. It demonstrates that the BBR step is the main source of variation in the ABCD-BIDS pipeline.

**Figure S4.1.**
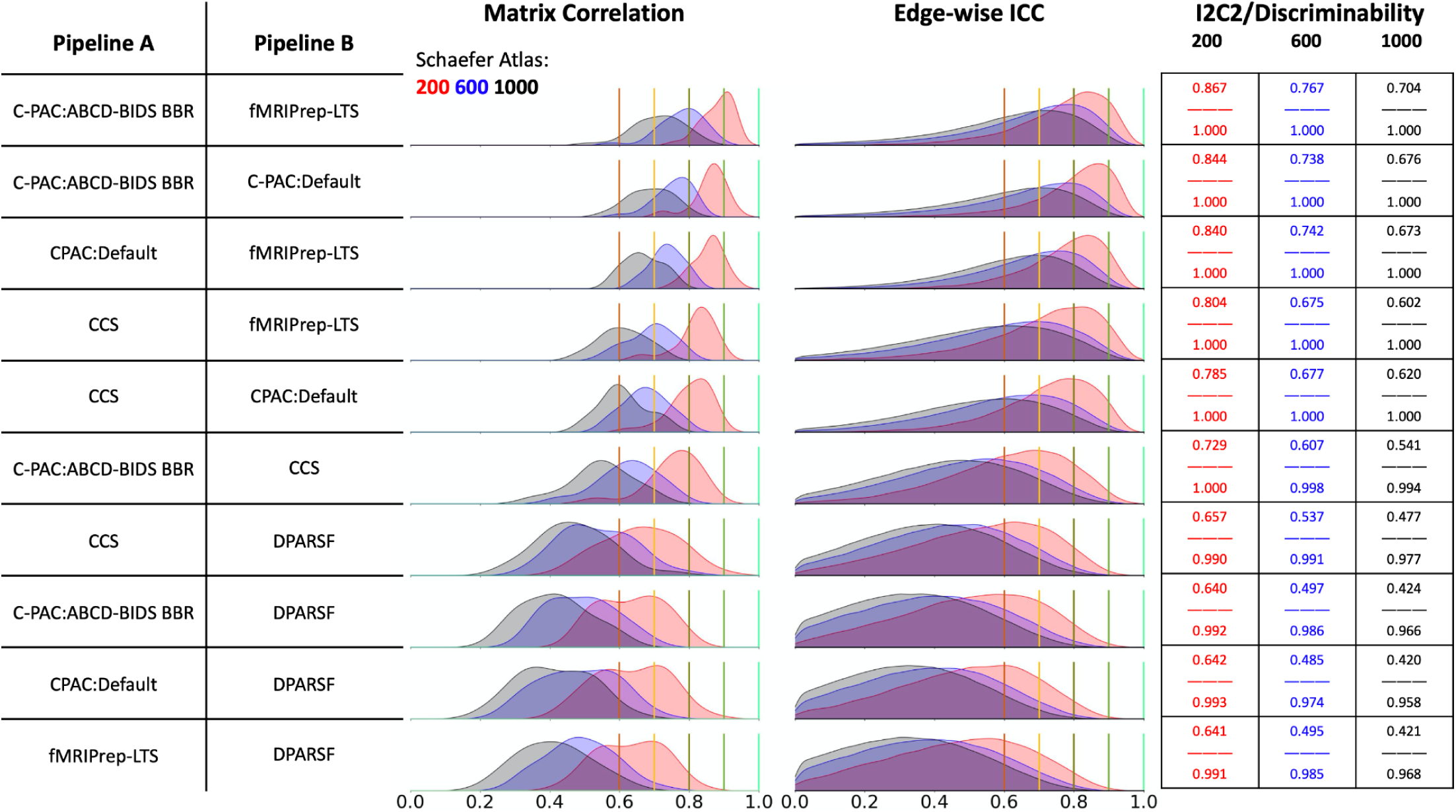
Modifying C-PAC:ABCD-BIDS to use boundary-based registration.

**S5: Inter-pipeline agreement with higher quality data**

A potential limitation in the implications of our presented findings is that they were generated using a functional imaging dataset that is behind the current state-of-the-art. While the selected dataset, HNU, was essential for our study given the large availability of test-retest measures, we replicated the cross-pipeline comparison using 29 low-motion subjects (mean FD Power: 0.118 ± 0.337 mm; mean FD Jenkinson: 0.068 ± 0.198 mm) and 29 high-motion subjects (mean FD Power: 0.681 ± 1.089 mm; mean FD Jenkinson: 0.389 ± 0.617 mm) from the HBN dataset, which uses a modern functional imaging sequence consistent with other major initiatives, such as the Adolescent Behavior and Cognitive Development dataset^25^. Shown in **Figure S5.1** and **Figure S5.2** below, we similarly found that inter-pipeline reliability was imperfect when looking across pipelines, even with high quality datasets. The reliability of measures improves compared to the HNU dataset, but still doesn’t meet accepted standards of inter-rater reliability, such as an ICC > 0.9.

**Figure S5.1.**
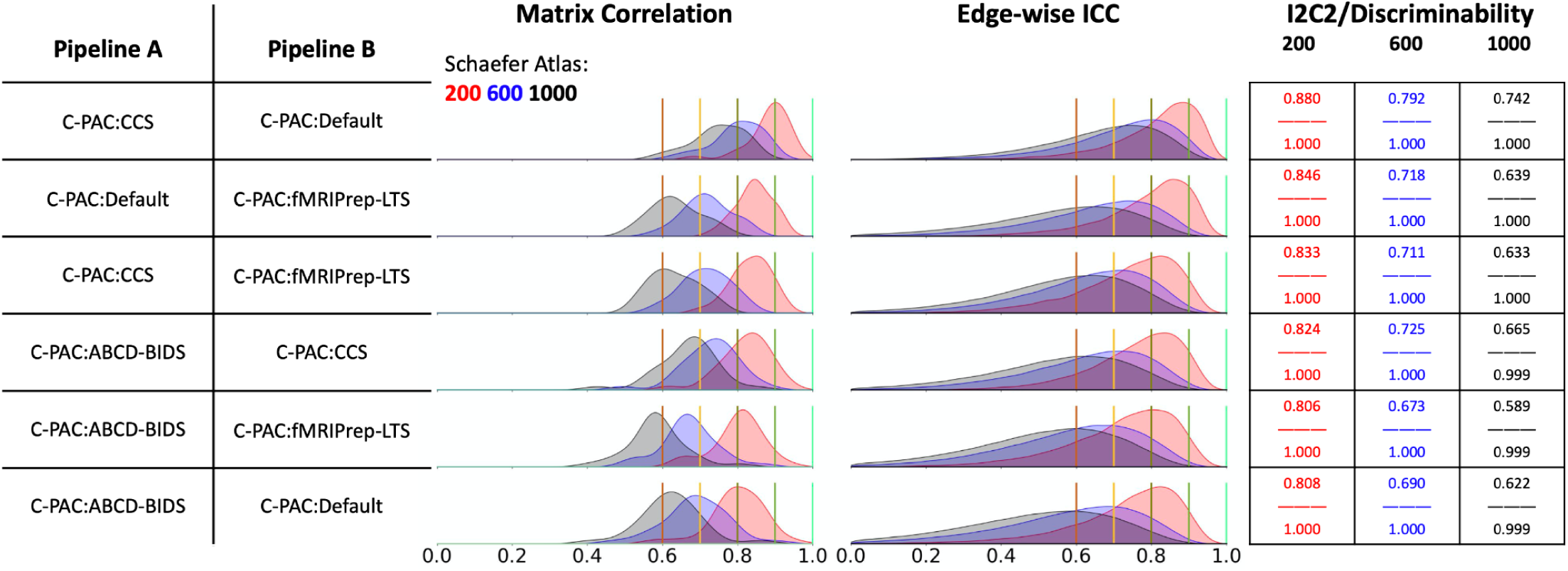
Inter-pipeline agreement with higher quality data (low-motion subjects).

**Figure S5.2.**
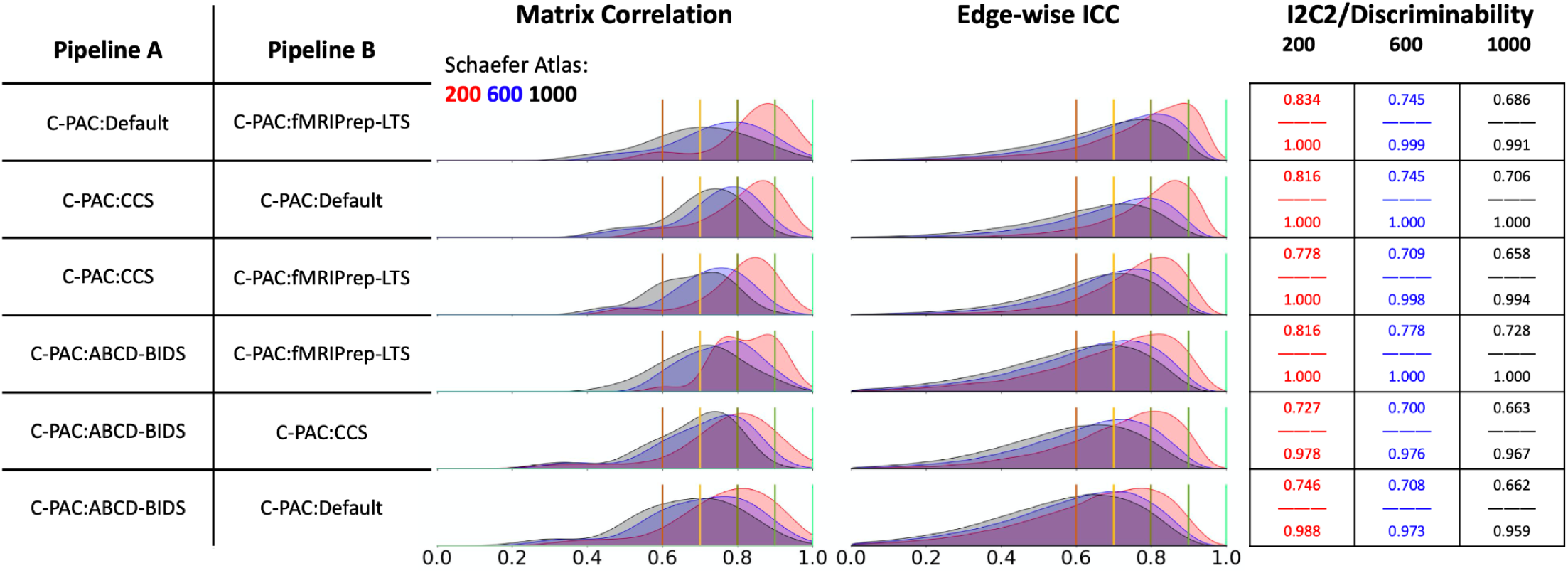
Inter-pipeline agreement with higher quality data (high-motion subjects).

**S6: Replication of intermediate results**

The figures below show the replication of intermediate derivatives across each of the three major harmonized packages. The intermediate derivatives include anatomical masks, white matter masks or white matter partial volume maps, functional masks, six motion parameters (rotation: rx, ry, rz; translation: tx, ty, tz), anatomical images and mean functional images in the MNI template space. The "Anat/Func-MNI pipeline” indicates the correlation between the pipeline and the standard template, e.g. “Anat-MNI ABCD-BIDS” in **Figure S6.1** refers to the correlation between the ABCD-BIDS pipeline output and the standard template; “Anat/Func-MNI” indicates the correlation between two pipelines, e.g. “Anat-MNI” in **Figure S6.1** refers to the correlation between the ABCD-BIDS pipeline output and the C-PAC:ABCD-BIDS pipeline output. Each column indicates a subject in the HNU dataset, and each row is an intermediate derivative. For each cell, the Pearson correlation between the derivatives across the two tools is shown. We recognize that the Pearson correlation may not be the most appropriate measure for some of the comparisons (e.g., aligned images), but it was used universally a) because it can be computed on all listed stages, and b) for consistency. When considering the anatomical images and mean functional images aligned to the MNI template, which are the primarily used derivatives in downstream analysis, we see that the lowest correlation is 0.84, while the majority of subjects have correlation values of above 0.97.

**Figure S6.1.**
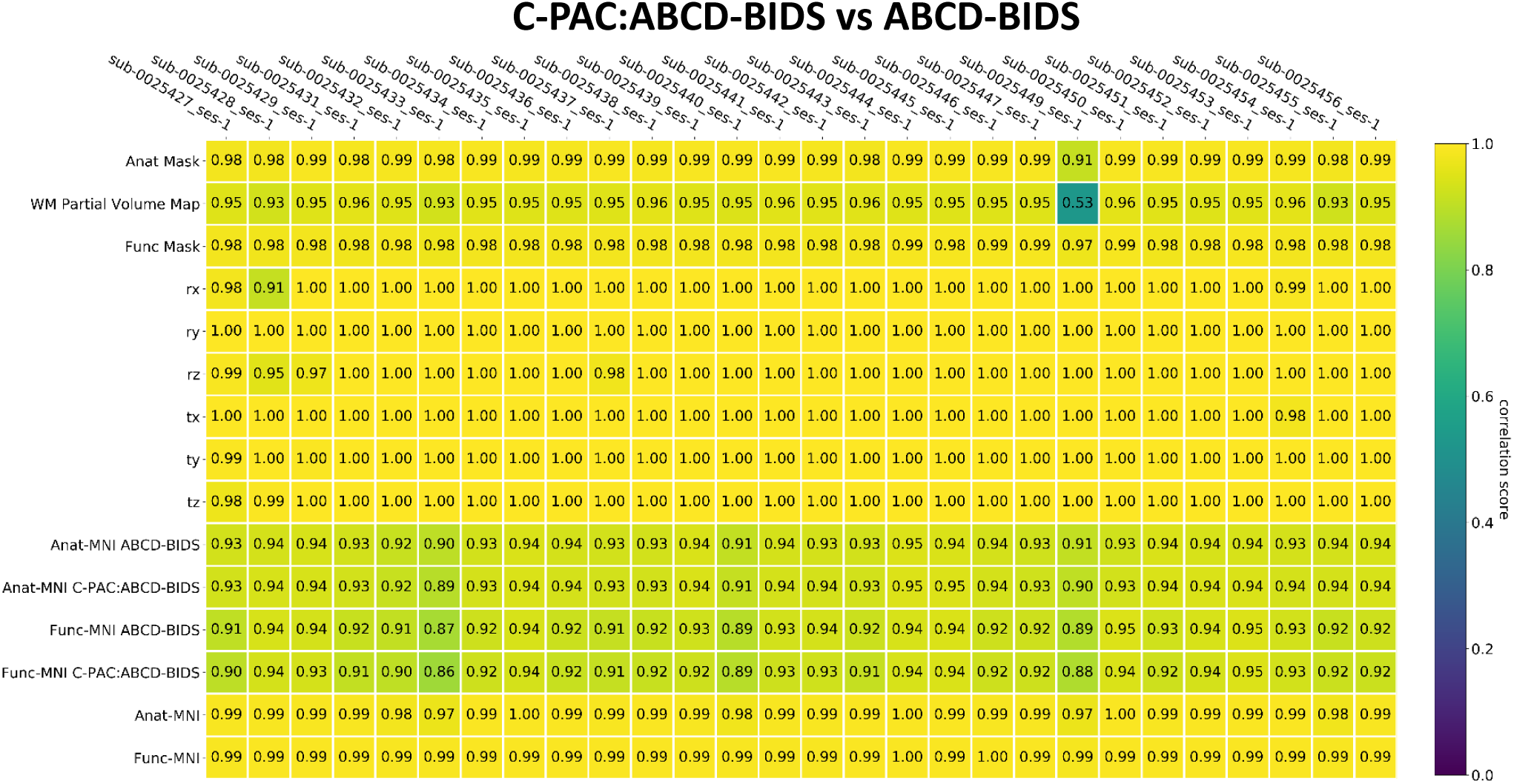
Reproducibility indices of intermediate derivatives for ABCD-BIDS.

**Figure S6.2.**
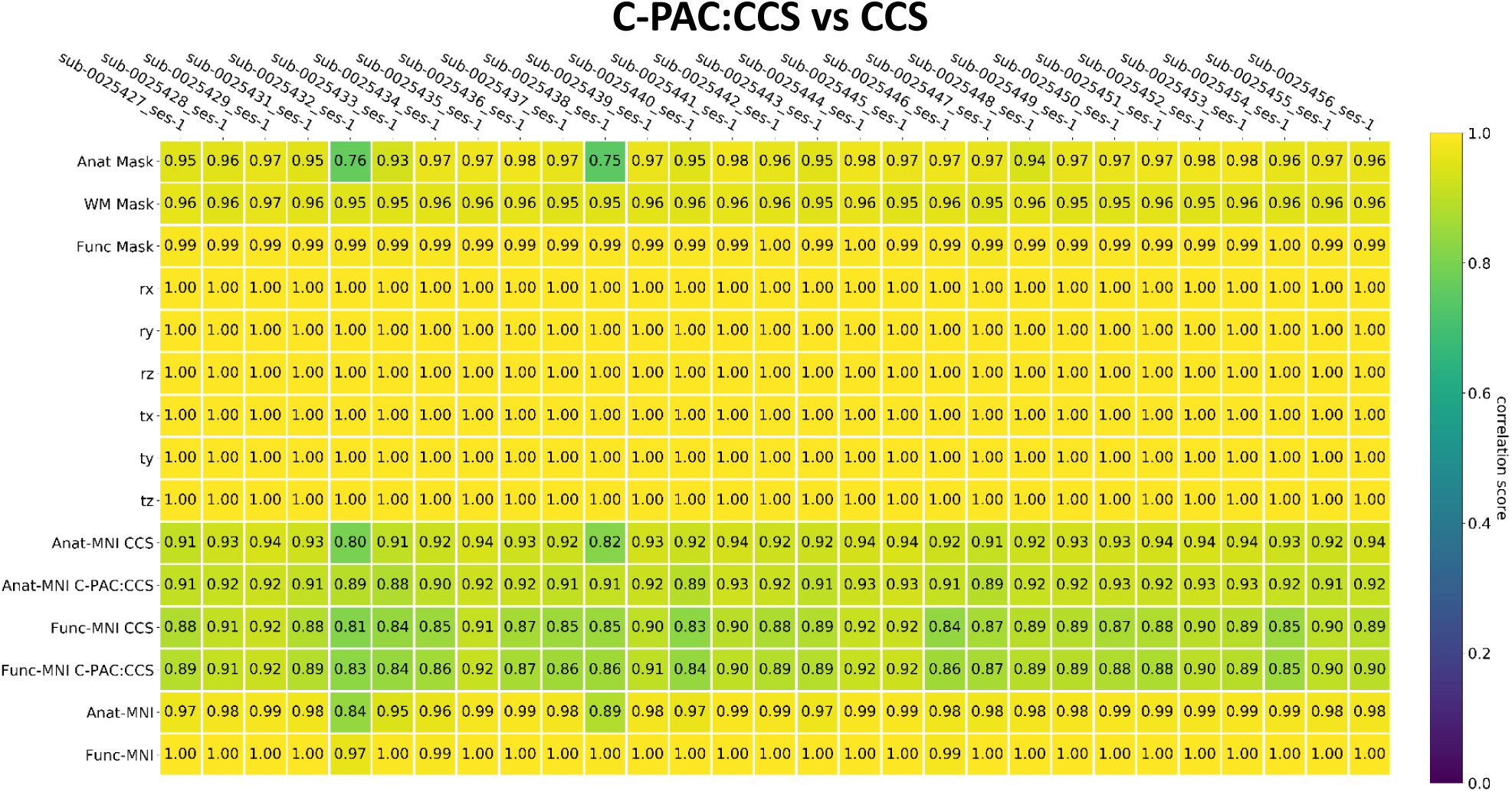
Reproducibility indices of intermediate derivatives for CCS.

**Figure S6.3.**
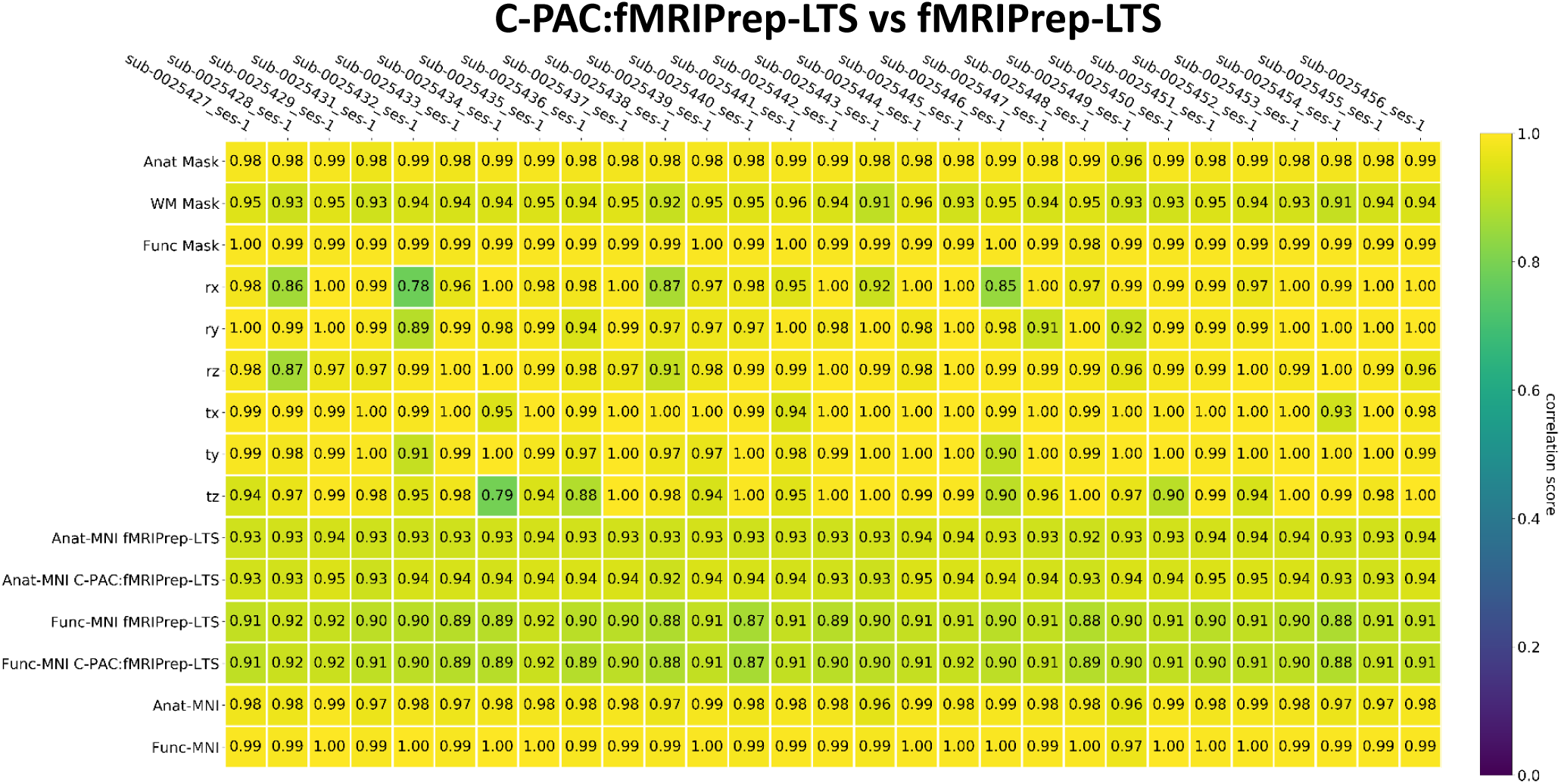
Reproducibility indices of intermediate derivatives for fMRIPrep-LTS.

**S7: Comparison of motion correction tools and references**

29 subjects with low motion (mean FD Power: 0.094 ± 0.190 mm; mean FD Jenkinson: 0.054 ± 0.107 mm) and 29 subjects with high motion (mean FD Power: 1.793 ± 3.414 mm; mean FD Jenkinson: 1.000 ± 1.887 mm) were selected from the HBN dataset. We evaluated the motion corrected outputs and the final functional time series in template space from two motion correction tools (AFNI 3dvolreg vs FSL MCFLIRT) and four motion correction references (mean volume, median volume, the first volume, the last volume) that are implemented in the C-PAC:Default pipeline. As shown in Figures S7.1 and S7.2, we observe greater variation in motion corrected time series across different references using FSL MCFLIRT than those using AFNI 3dvolreg, especially for the low-motion case. From **Figure S7.3**, we can see the moderate average correlation with large variance between FD results from AFNI 3dvolreg and those from FSL MCFLIRT when using the same reference. However, as shown in **Figure S7.4**, the final functional connectivity estimates are highly correlated regardless of motion correction implementations.

**Figure S7.1.**
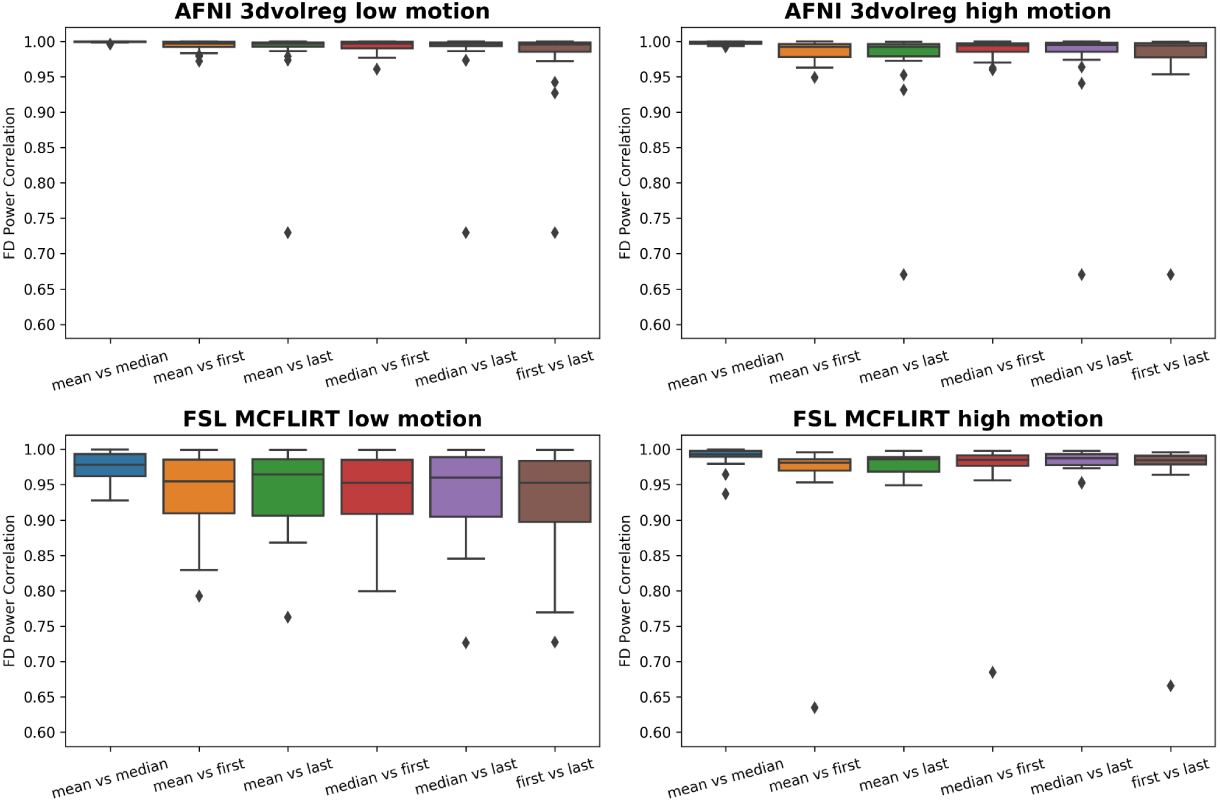
Comparison of motion correction tools and references. The top row shows AFNI 3dvolreg Power FD correlation results while the bottom row shows FSL MCFLIRT Power FD correlation results.

**Figure S7.2.**
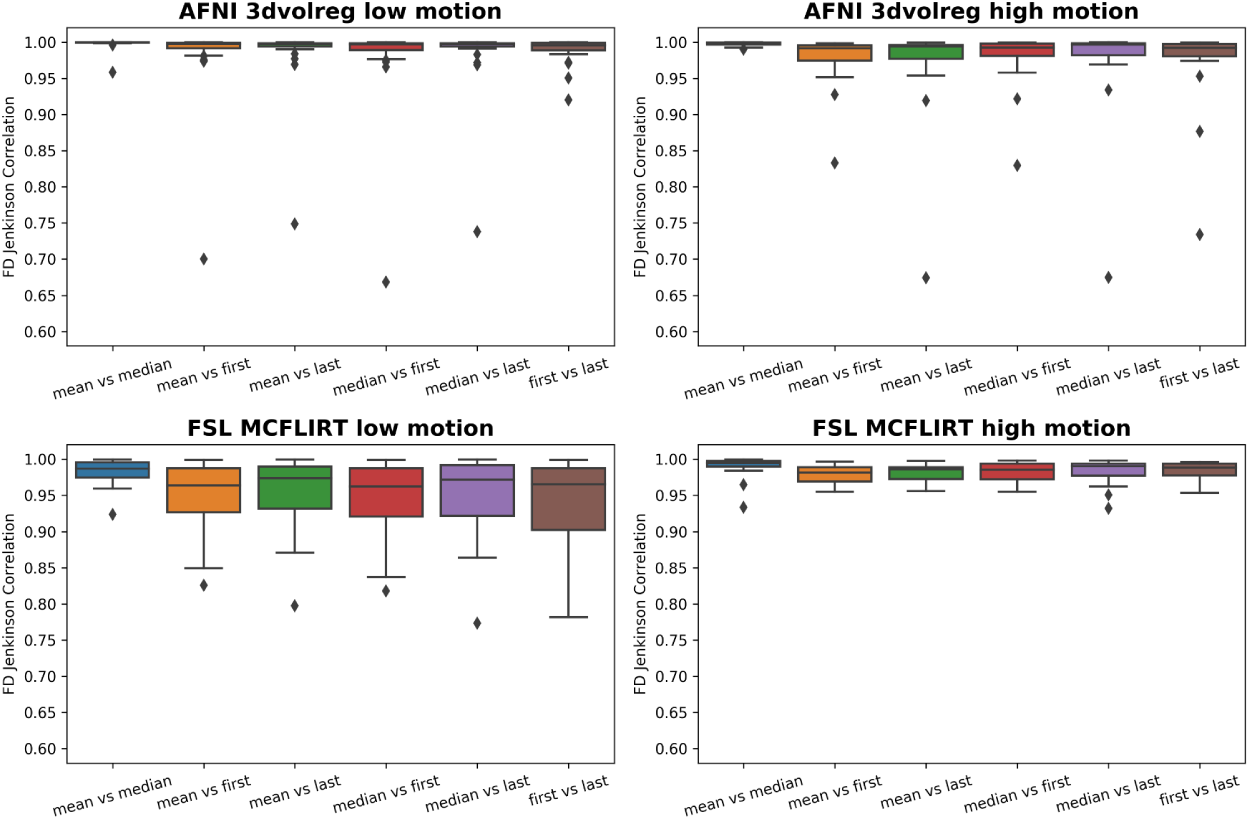
Comparison of motion correction tools and references. The top row shows AFNI 3dvolreg Jenkinson FD correlation results while the bottom row shows FSL MCFLIRT Jenkinson FD correlation results.

**Figure S7.3.**
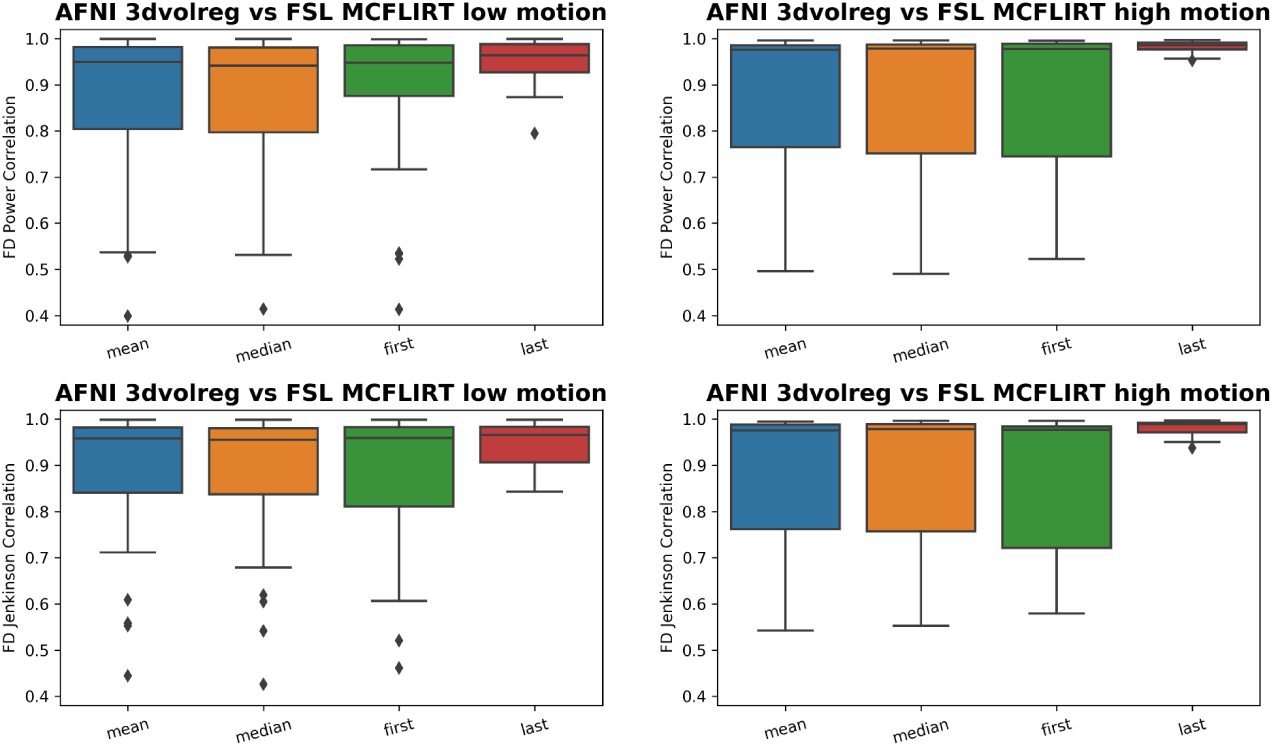
Comparison of motion correction tools and references. The top row shows AFNI 3dvolreg vs FSL MCFLIRT Power FD correlation results while the bottom row shows Jenkinson FD correlation results.

**Figure S7.4.**
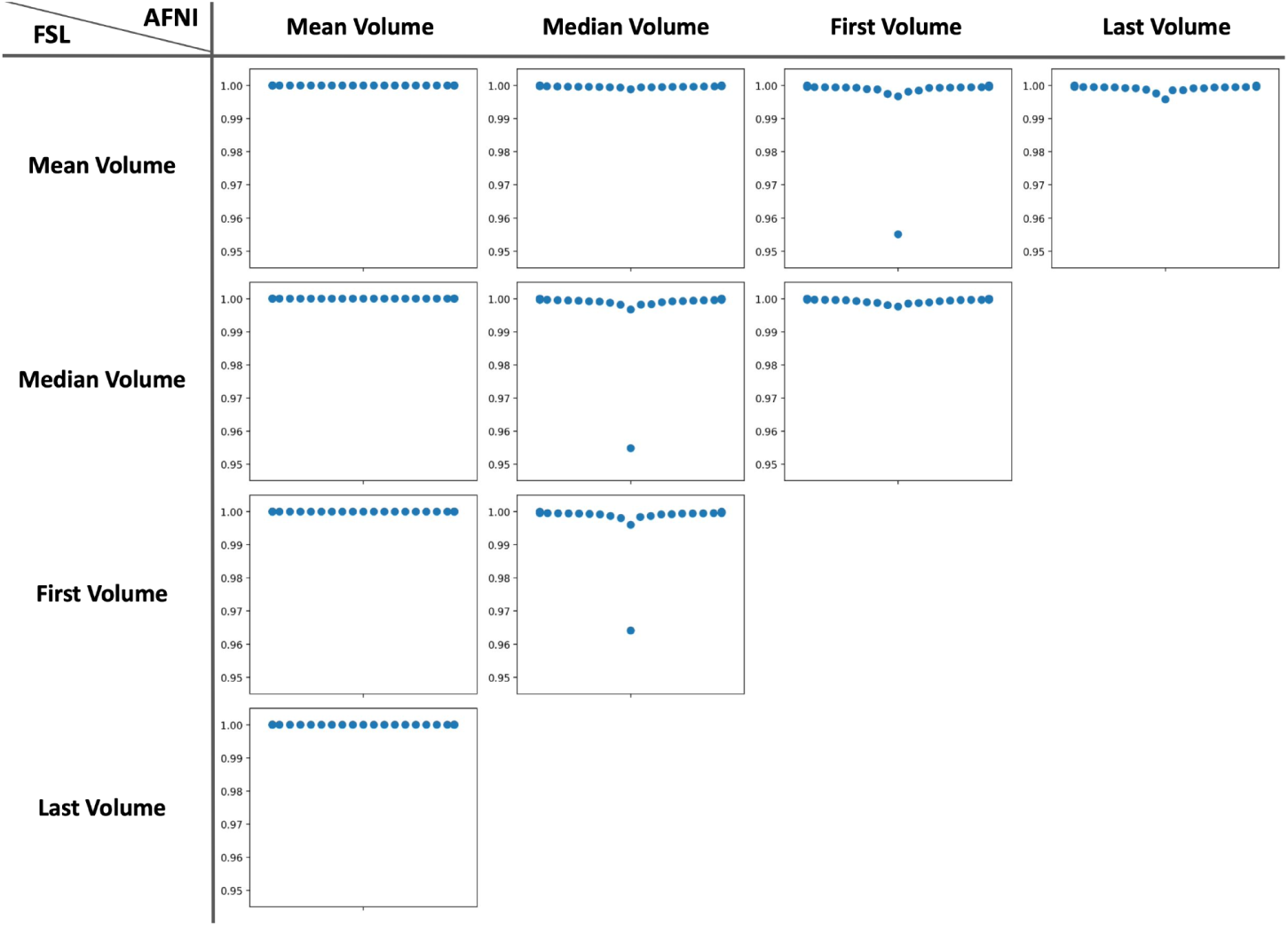
Comparison of motion correction tools and references. Pearson correlation of functional connectivity matrices using Schaefer 200 unit parcellation. Each dot indicates Pearson correlation for one subject.

**S8: Validation of impact of MNI template version and write-out resolution**

We computed spatial correlation of voxel-wise time series to further validate the intra-pipeline agreement when different MNI template versions and write-out resolution were used. We resampled the functional time series to a common space (defined by the MNI2006 template) with matched resolution using AFNI 3dresample, because the 2009 template has a different number of voxels from the 2001 and 2006 templates. We performed voxel-wise spatial correlation of the time series using AFNI 3dTcorrelate, and then calculated the mean, standard deviation, and the 5, 50, and 95^th^ percentiles of spatial correlation across configurations using AFNI 3dBrickStat. As shown in Table S8.1, the spatial correlation of voxel-wise time series across the 2001 and 2006 versions of the MNI template show high correspondence in both write-out settings, while results from the 2009 template show significantly lower correspondence. To avoid inconsistency in data manipulations and confounding sources of bias in this analysis, no comparisons were made with unmatched write-out resolutions (i.e. native versus 2mm). The result here aligns with findings shown in Figure 6.

**Table S8.1.**
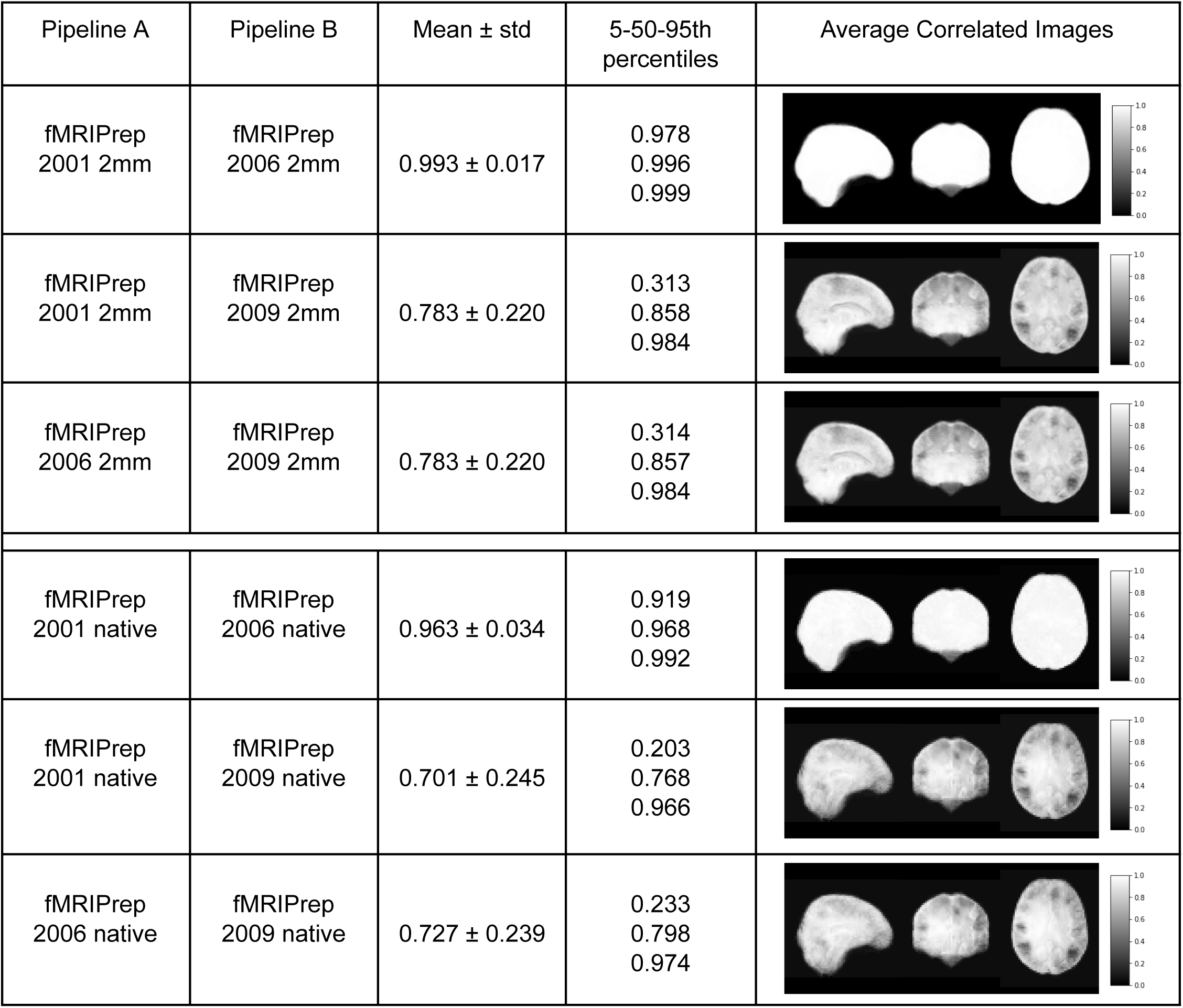
Voxel-wise time series correlation of different MNI template versions and write-out resolution.

| Pipeline A | Pipeline B | Mean ± std | 5-50-95th percentiles | Average Correlated Images |
| --- | --- | --- | --- | --- |
| fMRIPrep<br>2001 2mm | fMRIPrep<br>2006 2mm | $0.993 \pm 0.017$ | 0.978<br>0.996<br>0.999 | |
| fMRIPrep<br>2001 2mm | fMRIPrep<br>2009 2mm | $0.783 \pm 0.220$ | 0.313<br>0.858<br>0.984 | |
| fMRIPrep<br>2006 2mm | fMRIPrep<br>2009 2mm | $0.783 \pm 0.220$ | 0.314<br>0.857<br>0.984 | |
| fMRIPrep<br>2001 native | fMRIPrep<br>2006 native | $0.963 \pm 0.034$ | 0.919<br>0.968<br>0.992 | |
| fMRIPrep<br>2001 native | fMRIPrep<br>2009 native | $0.701 \pm 0.245$ | 0.203<br>0.768<br>0.966 | |
| fMRIPrep<br>2006 native | fMRIPrep<br>2009 native | $0.727 \pm 0.239$ | 0.233<br>0.798<br>0.974 | |

## References

1. Shehzad, Z. et al. The resting brain: unconstrained yet reliable. Cereb. Cortex 19, 2209–2229 (2009).

2. Zuo, X.-N. et al. The oscillating brain: complex and reliable. Neuroimage 49, 1432–1445 (2010).

3. Bennett, C. M. & Miller, M. B. How reliable are the results from functional magnetic resonance imaging? Ann. N. Y. Acad. Sci. 1191, 133–155 (2010).

4. Zuo, X.-N., Xu, T. & Milham, M. P. Harnessing reliability for neuroscience research. Nat Hum Behav 3, 768–771 (2019).

5. Kraemer, H. C. The reliability of clinical diagnoses: state of the art. Annu. Rev. Clin. Psychol. 10, 111–130 (2014).

6. Button, K. S. et al. Power failure: why small sample size undermines the reliability of neuroscience. Nat. Rev. Neurosci. 14, 365–376 (2013).

7. Ioannidis, J. P. A. Why most published research findings are false. PLoS Med. 2, e124 (2005).

8. Noble, S., Scheinost, D. & Constable, R. T. A decade of test-retest reliability of functional connectivity: A systematic review and meta-analysis. Neuroimage 203, 116157 (2019).

9. Zuo, X.-N. & Xing, X.-X. Test-retest reliabilities of resting-state FMRI measurements in human brain functional connectomics: a systems neuroscience perspective. Neurosci. Biobehav. Rev. 45, 100–118 (2014).

10. Cho, J. W., Korchmaros, A., Vogelstein, J. T., Milham, M. P. & Xu, T. Impact of concatenating fMRI data on reliability for functional connectomics. Neuroimage 226, 117549 (2021).

11. Lynch, C. J. et al. Rapid Precision Functional Mapping of Individuals Using Multi-Echo fMRI. Cell Rep. 33, 108540 (2020).

12. Nikolaidis, A. et al. Bagging improves reproducibility of functional parcellation of the human brain. Neuroimage 214, 116678 (2020).

13. Yoo, K. et al. Multivariate approaches improve the reliability and validity of functional connectivity and prediction of individual behaviors. Neuroimage 197, 212–223 (2019).

14. Elliott, M. L. et al. What Is the Test-Retest Reliability of Common Task-Functional MRI Measures? New Empirical Evidence and a Meta-Analysis. Psychol. Sci. 31, 792–806 (2020).

15. Palumbo, L. et al. Evaluation of the intra- and inter-method agreement of brain MRI segmentation software packages: A comparison between SPM12 and FreeSurfer v6.0. Phys. Med. 64, 261–272 (2019).

16. Oakes, T. R. et al. Comparison of fMRI motion correction software tools. Neuroimage 28, 529–543 (2005).

17. Klein, A. et al. Evaluation of 14 nonlinear deformation algorithms applied to human brain MRI registration. Neuroimage 46, 786–802 (2009).

18. Dickie, E., Hodge, S., Craddock, R., Poline, J.-B. & Kennedy, D. Tools matter: Comparison of two surface analysis tools applied to the ABIDE dataset. Res. Ideas Outcomes 3, e13726 (2017).

19. Bhagwat, N. et al. Understanding the impact of preprocessing pipelines on neuroimaging cortical surface analyses. Gigascience 10, (2021).

20. Carp, J. On the plurality of (methodological) worlds: estimating the analytic flexibility of FMRI experiments. Front. Neurosci. 6, 149 (2012).

21. Pauli, R. et al. Exploring fMRI Results Space: 31 Variants of an fMRI Analysis in AFNI, FSL, and SPM. Front. Neuroinform. 10, 24 (2016).

22. Bowring, A., Maumet, C. & Nichols, T. E. Exploring the impact of analysis software on task fMRI results. Hum. Brain Mapp. 40, 3362–3384 (2019).

23. Bowring, A., Nichols, T. E. & Maumet, C. Isolating the sources of pipeline-variability in group-level task-fMRI results. Hum. Brain Mapp. (2021) doi:10.1002/hbm.25713.

24. Botvinik-Nezer, R. et al. Variability in the analysis of a single neuroimaging dataset by many teams. Nature 582, 84–88 (2020).

25. Feczko, E., Conan, G., Marek, S. & Tervo-Clemmens, B. Adolescent Brain Cognitive Development (ABCD) Community MRI Collection and Utilities. bioRxiv (2021).

26. Xu, T., Yang, Z., Jiang, L., Xing, X.-X. & Zuo, X.-N. A Connectome Computation System for discovery science of brain. Sci Bull. Fac. Agric. Kyushu Univ. 60, 86–95 (2015).

27. Craddock, C. et al. Towards automated analysis of connectomes: The configurable pipeline for the analysis of connectomes (c-pac). Front. Neuroinform. 42, 10–3389 (2013).

28. Chao-Gan, Y. & Yu-Feng, Z. DPARSF: A MATLAB toolbox for ‘pipeline’ data analysis of resting-State fMRI. Front. Syst. Neurosci. 4, 13 (2010).

29. Esteban, O. et al. fMRIPrep: a robust preprocessing pipeline for functional MRI. Nat. Methods 16, 111–116 (2019).

30. Murphy, K. & Fox, M. D. Towards a consensus regarding global signal regression for resting state functional connectivity MRI. Neuroimage 154, 169–173 (2017).

31. Zuo, X.-N. et al. An open science resource for establishing reliability and reproducibility in functional connectomics. Sci Data 1, 140049 (2014).

32. Shou, H. et al. Quantifying the reliability of image replication studies: the image intraclass correlation coefficient (I2C2). Cogn. Affect. Behav. Neurosci. 13, 714–724 (2013).

33. Bridgeford, E. W. et al. Eliminating accidental deviations to minimize generalization error and maximize replicability: Applications in connectomics and genomics. PLoS Comput. Biol. 17, e1009279 (2021).

34. Schaefer, A. et al. Local-Global Parcellation of the Human Cerebral Cortex from Intrinsic Functional Connectivity MRI. Cereb. Cortex 28, 3095–3114 (2018).

35. Glasser, M. F. et al. The minimal preprocessing pipelines for the Human Connectome Project. Neuroimage 80, 105–124 (2013).

36. Greve, D. N. & Fischl, B. Accurate and robust brain image alignment using boundary-based registration. Neuroimage 48, 63–72 (2009).

37. Alexander, L. M. et al. An open resource for transdiagnostic research in pediatric mental health and learning disorders. Sci Data 4, 170181 (2017).

38. Koo, T. K. & Li, M. Y. A Guideline of Selecting and Reporting Intraclass Correlation Coefficients for Reliability Research. Journal of Chiropractic Medicine vol. 15 155–163 Preprint at 10.1016/j.jcm.2016.02.012 (2016).

39. Dong, Y., Ifrim, G., Mladenić, D., Saunders, C. & Van Hoecke, S. Machine Learning and Knowledge Discovery in Databases. Applied Data Science and Demo Track: European Conference, ECML PKDD 2020, Ghent, Belgium, September 14–18, 2020, Proceedings, Part V. (Springer Nature, 2021).

40. Birn, R. M. et al. The effect of scan length on the reliability of resting-state fMRI connectivity estimates. Neuroimage 83, 550–558 (2013).

41. Liu, T. T., Nalci, A. & Falahpour, M. The global signal in fMRI: Nuisance or Information? Neuroimage 150, (2017).

42. Ciric, R. et al. Benchmarking of participant-level confound regression strategies for the control of motion artifact in studies of functional connectivity. Neuroimage 154, 174–187 (2017).

43. Buades, A., Coll, B. & Morel, J.-M. Non-Local Means Denoising. Image process. line 1, 208–212 (2011).

44. Tustison, N. J. et al. N4ITK: improved N3 bias correction. IEEE Trans. Med. Imaging 29, 1310–1320 (2010).

45. Ciric, R., Lorenz, R., Thompson, W. H. & Goncalves, M. TemplateFlow: a community archive of imaging templates and atlases for improved consistency in neuroimaging. bioRxiv (2021).

46. Fonov, V. S., Evans, A. C., McKinstry, R. C., Almli, C. R. & Collins, D. L. Unbiased nonlinear average age-appropriate brain templates from birth to adulthood. Neuroimage Supplement 1, S102 (2009).

47. Grabner, G. et al. Symmetric atlasing and model based segmentation: an application to the hippocampus in older adults. Med. Image Comput. Comput. Assist. Interv. 9, 58–66 (2006).

48. Mazziotta, J. et al. A probabilistic atlas and reference system for the human brain: International Consortium for Brain Mapping (ICBM). Philos. Trans. R. Soc. Lond. B Biol. Sci. 356, 1293–1322 (2001).

49. Wu, J. et al. Accurate nonlinear mapping between MNI volumetric and FreeSurfer surface coordinate systems. Hum. Brain Mapp. 39, 3793–3808 (2018).

50. Uddin, L. Q. Mixed Signals: On Separating Brain Signal from Noise. Trends in cognitive sciences vol. 21 405–406 (2017).

51. Murphy, K., Birn, R. M., Handwerker, D. A., Jones, T. B. & Bandettini, P. A. The impact of global signal regression on resting state correlations: are anti-correlated networks introduced? Neuroimage 44, 893–905 (2009).

52. Xu, H. et al. Impact of global signal regression on characterizing dynamic functional connectivity and brain states. Neuroimage 173, 127–145 (2018).

53. Gordon, E. M. et al. Precision Functional Mapping of Individual Human Brains. Neuron 95, 791–807.e7 (2017).

54. Di Martino, A. et al. Enhancing studies of the connectome in autism using the autism brain imaging data exchange II. Sci Data 4, 170010 (2017).

55. Casey, B. J. et al. The Adolescent Brain Cognitive Development (ABCD) study: Imaging acquisition across 21 sites. Dev. Cogn. Neurosci. 32, 43–54 (2018).

56. Doshi, J. et al. MUSE: MUlti-atlas region Segmentation utilizing Ensembles of registration algorithms and parameters, and locally optimal atlas selection. NeuroImage vol. 127 186–195 Preprint at 10.1016/j.neuroimage.2015.11.073 (2016).

57. Wu, D. et al. Resource atlases for multi-atlas brain segmentations with multiple ontology levels based on T1-weighted MRI. Neuroimage 125, 120–130 (2016).

58. Kiar, G., Chatelain, Y., Salari, A., Evans, A. C. & Glatard, T. Data Augmentation Through Monte Carlo Arithmetic Leads to More Generalizable Classification in Connectomics. Neurons, Behaviour, Data, and Theory (2021) doi:10.51628/001c.28328.

59. Kiar, G. et al. Numerical Uncertainty in Analytical Pipelines Lead to Impactful Variability in Brain Networks. PLoS One 2020.10.15.341495 (2021).

60. Mehta, K., et al. XCP-D: A Robust Pipeline for the post-processing of fMRI data. bioRxiv (2023) doi:10.1101/2023.11.20.567926.

61. Bujang & Baharum. A simplified guide to determination of sample size requirements for estimating the value of intraclass correlation coefficient: a review. Arch. orofac. sci.

62. Smith, S. M. et al. Advances in functional and structural MR image analysis and implementation as FSL. Neuroimage 23 Suppl 1, S208–19 (2004).

63. Jenkinson, M., Bannister, P., Brady, M. & Smith, S. Improved optimization for the robust and accurate linear registration and motion correction of brain images. Neuroimage 17, 825–841 (2002).

64. Cox, R. W. AFNI: Software for Analysis and Visualization of Functional Magnetic Resonance Neuroimages. Comput. Biomed. Res. 29, 162–173 (1996).

65. Zhang, Y., Brady, M. & Smith, S. Segmentation of brain MR images through a hidden Markov random field model and the expectation-maximization algorithm. IEEE Transactions on Medical Imaging vol. 20 45–57 Preprint at 10.1109/42.906424 (2001).

66. Avants, B. B., Tustison, N., Song, G. & Others. Advanced normalization tools (ANTS). Insight J. 2, 1–35 (2009).

67. Fischl, B. FreeSurfer. Neuroimage 62, 774–781 (2012).

68. Jenkinson, M. & Smith, S. A global optimisation method for robust affine registration of brain images. Med. Image Anal. 5, 143–156 (2001).

69. Berger, V. W. & Zhou, Y. Kolmogorov–Smirnov Test: Overview. Wiley StatsRef: Statistics Reference Online Preprint at 10.1002/9781118445112.stat06558 (2014).

70. Nachar, N. The Mann-Whitney U: A test for assessing whether two independent samples come from the same distribution. Tutor. Quant. Methods Psychol. 4, 13–20 (2008).

